# Flightless I and LRRFIP work together to regulate lateral growth of the sarcomeres in *Drosophila*

**DOI:** 10.64898/2026.06.02.729301

**Authors:** Péter Görög, Krisztina Tóth, Dávid Farkas, Balázs Vedelek, Tamás F. Polgár, Anna Zsuzsanna Tihanyi, Péter Bíró, Tibor Novák, Johannes Salomonsson, Kristina Djinovic Carugo, Aladár Pettkó-Szandtner, Zsuzsanna Darula, Miklós Erdélyi, Szilárd Szikora, József Mihály

## Abstract

Myofibrils are structurally highly conserved elements of the cardiac and striated muscles. The striking commonalities in their basic organization principles emerge during myofibrillogenesis when the initially formed premyofibrils grow in length and width to attain their final dimensions, characteristic of each muscle type. Although the molecular composition of the myofibrils is well known, the mechanisms governing their growth remain poorly understood, particularly those driving peripheral thickening. Here, we show that two cardiac disease associated proteins, Flightless I and LRRFIP, are required for lateral integration of the myofilaments that is essential for circumferential myofibril growth in the *Drosophila* flight muscle. Genetic and biochemical analysis reveal that these proteins form multimeric complexes enriched at the barbed end of the actin filaments. Furthermore, we found that the Flightless I/LRRFIP complex acts in a formin dependent manner. Together, our findings demonstrate that Flightless I and LRRFIP cooperate to promote the spatially controlled integration of peripheral actin filaments at the Z-disc, uncovering a key step in myofibril maturation.

## INTRODUCTION

Despite the great diversity in the physiological demands, and therefore in the size and physical properties of the various muscles found in nature, all striated and cardiac muscles share a similar structural organization, comprising myofibers typically made up of dozens to hundreds of myofibrils. The thread-like myofibrils consist of sarcomeres, serially repeated units bordered by Z-discs, known as the smallest contractile elements of the muscles. Whereas the precise details of the first steps of myofibril formation are incompletely understood, the end of the initial phase is marked by the formation of myofibril precursors, called premyofibrils. These structures are composed of premature sarcomeres, primarily built up from the major sarcomeric components, including proteins of the thin, thick and elastic filaments. Nevertheless, to attain their highly stereotypical mature size, the premyofibrillar sarcomeres must undergo a three-dimensional expansion during the subsequent phases of myofibrillogenesis, resulting in a significant increase both in their length and width (Sanger et al., 2017; Szikora et al., 2022).

The first insights into the mechanisms of this growth process were gained in GFP-labeled actin incorporation studies in cardiomyocytes, revealing that the sarcomeric actin filaments mainly elongate from their pointed end (located at the edges of the H-zone) (Fig. 1a) (Littlefield et al., 2001), and that lateral/circumferential growth of the premyofibrillar sarcomeres occurs by actin incorporation at the periphery of the sarcomeres (Sanger et al., 2005; Shwartz et al., 2016). Genetic perturbation of members of the Leimodin (Lmod) and formin protein families (Campellone and Welch, 2010), exhibiting actin assembly activities (Chesarone et al., 2010; Fowler and Dominguez, 2017), and that of the pointed end capping Tropomodulin (Tmod) protein, gave rise to altered sarcomere length (Chereau et al., 2008; Littlefield et al., 2001; Mardahl-Dumesnil and Fowler, 2001; Tsukada et al., 2010). Similarly, the loss or overexpression of the *Drosophila* actin binding protein, SALS (Sarcomere Length Short), also results in sarcomere length alterations (Bai et al., 2007; Farkas et al., 2024). Whereas, with the exception of Tmod, all these proteins exhibit Z-disc (i.e. barbed end) and H-zone (i.e. pointed end) accumulation as well, in line with the GFP::actin incorporation studies, the prevailing view is that the sarcomeric actin filaments primarily elongate from or at their pointed end (Szikora et al., 2022). However, due to their complex muscle phenotypes and the partly controversial *in vitro* data, it remained poorly understood how these factors synergize to promote filament elongation from the pointed end. Likewise, although the ends of the thin filaments are in perfect register with each other at the edges of the Z-discs and at the borders of the H-zone, the mechanisms controlling filament length, and in particular, that of filament end alignment, are also unknown.

**Figure 1.**
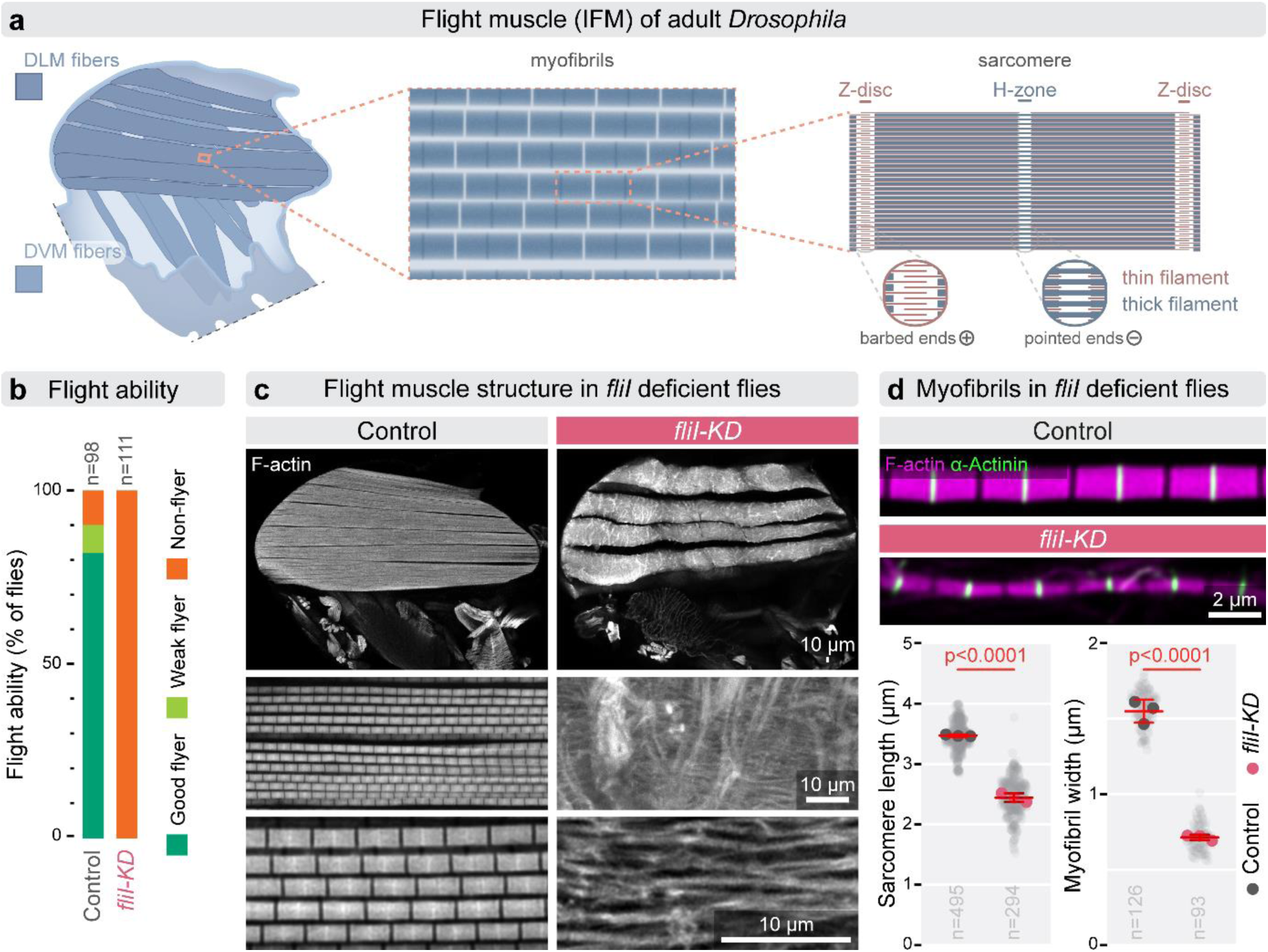
The knockdown of FliI impairs myofibril growth in the *Drosophila* IFM. **(a)** Schematic of the indirect flight muscles (IFMs), which consist of the dorsal longitudinal muscles (DLM, dark blue) and dorso-ventral muscles (DVM, light blue). Myofibrils are composed of regularly arranged sarcomeres with defined length and diameter. Z-discs, marking sarcomere boundaries, and the central H-zone are indicated, with thin filament orientation highlighted. **(b)** Flight assay showing complete loss of flight in *fliI-KD* (*mef2-Gal4/UAS-TRIP.JF02720*) flies at ∼24 hours AE, compared with ∼90% flight ability in controls (*w^1118^*). **(c)** Low- and high-magnification images of sagittal thoracic sections reveal DLM fiber structure. In *fliI-KD* animals, large myofibrillar aggregates are visible on the surface of otherwise normally sized fibers. At higher resolution, individual myofibrils appear thinner, disorganized, and exhibit severely disrupted sarcomeric structure with loss of the regular actin pattern. F-actin is visualized by phalloidin staining (gray). **(d)** Top panels show isolated myofibrils from control and *fliI-KD* IFMs, demonstrating reduced sarcomere length and myofibril diameter in knockdown animals. The graph below presents morphometric quantification, confirming significant reductions in both parameters (p<0.0001). Statistical significance was assessed using unpaired t-tests. Light gray dots represent individual measurements, while larger dots indicate mean values from independent experiments. Error bars show mean of independent experiments ± s.d., and n denotes the number of independent measurements. Raw data are provided in Fig1SourceData.

Regarding the mechanisms of peripheral sarcomere growth, analysis of members of the Zasp protein family revealed that in *Drosophila* indirect flight muscles (IFM) the balance of the short (blocking) and long (growing) isoforms of Zasp52 is crucial to set the width of the myofibrils (Gonzalez-Morales et al., 2019). Because the imbalance of these isoforms results in enlarged or reduced Z-discs (and sometimes in actin aggregate formation), it was proposed that peripheral/radial sarcomere growth is initiated at the Z-disc. In vertebrates the Zasp proteins are known as Alp/Enigma family (Fisher and Schock, 2022), and in mouse models it was shown that in the absence of these proteins the sarcomeres look considerably smaller in diameter (Cheng et al., 2010; Mu et al., 2015), suggesting an evolutionary conserved role for the Zasp/Enigma proteins in setting the width of the myofibrils. Besides these findings, several studies described *Drosophila* mutants, such as *DAAM* and *fhos* (encoding formin family actin assembly factors) (Molnar et al., 2014; Shwartz et al., 2016), *sals* (Bai et al., 2007; Farkas et al., 2024), *sls* (Sallimus) (Orfanos et al., 2015), *Lasp* (Fernandes and Schock, 2014) and *fln* (*flightin*) (Chakravorty et al., 2017), where myofibril width was affected, but the mechanisms of circumferential enlargement of the sarcomeres were beyond the focus of those investigations. One additional gene belonging to this group is *flightless I* (*fliI*), encoding a highly conserved member of the gelsolin protein superfamily, consisting of a leucine-rich repeat region (LRR) at its N-terminus and 6 gelsolin homology (GH) domains at its C-terminus (Campbell et al., 1993). FliI has originally been identified with hypomorphic mutations impairing flight muscle development (Campbell et al., 1993; Deak et al., 1982; Miklos and De Couet, 1990). It was subsequently shown that *fliI* is required for cellularization and gastrulation in *Drosophila* embryos (Miklos and De Couet, 1990; Perrimon et al., 1989; Straub et al., 1996), and analogously, it is essential during embryonic development in mice and zebrafish (Campbell et al., 2002; Naganawa and Hirata, 2011). In addition, it was also revealed that FliI participates in numerous cellular and signaling processes, including proliferation, differentiation, cell migration and inflammation (Cowin et al., 2007; Kopecki et al., 2009; Nag et al., 2013; Strudwick and Cowin, 2020). Importantly, a further set of studies established that the loss or partial loss of *fliI* cause cardiomyopathy in both human and mice (Al-Hassnan et al., 2020; Karczewski et al., 2020; Kuwabara et al., 2023; Lipov et al., 2023; Ruijmbeek et al., 2023), and skeletal muscle fiber disorganization in zebrafish (Granato et al., 1996; Naganawa and Hirata, 2011). Despite these advances in linking FliI to muscle development and cardiac disease, the muscle phenotypes reported are clearly complex in every model systems examined, leading to (at least partly) conflicting conclusions as to the cellular and molecular function of FliI, i.e. a role in the regulation of sarcomeric actin dynamics and length (Kuwabara et al., 2023) or an effect on cell adhesion and heart trabeculation (Ruijmbeek et al., 2023), and an actin severing based mechanism (Deng et al., 2021) were all proposed as potential mode of actions. In addition, although FliI has been shown to localize to actin-based structures *in vivo* (Davy et al., 2001) and *in vitro* (Goshima et al., 1999; Liu and Yin, 1998; Mohammad et al., 2012; Pintér et al., 2020) via its GH domains, its role in thin filament (sarcomeric actin) regulation remained largely unclear, motivating us to revisit the role of FliI during muscle development.

In this study, we show that *Drosophila* FliI is required for multiple aspects of myofibrillogenesis, in particular, circumferential growth of the sarcomeres. FliI exhibits a barbed end association at the edges of the Z-disc, and accordingly, the loss of *fliI* impairs Z-disc formation. Our transmission electron microscopy (TEM) analysis revealed that FliI is not required for premyofibril formation nor for construction of the myofilaments that ensure myofibril growth, instead it is essential for peripheral integration of the myofilaments. In addition, we also show that FliI forms a complex with *Drosophila* LRRFIP (Leucine rich repeat FliI interacting protein), and they act in concert to promote radial sarcomere growth in a formin dependent manner.

## RESULTS

### FliI regulates sarcomere size and peripheral myofilament integration

To gain further insights into the muscle function of *Drosophila* FliI, we focused on the IFM (Fig. 1a) by characterizing two, adult viable, hypomorphic alleles, *fliI^3^* and *fliI^sdby^* (carrying point mutations in the GH2 and GH6 domains, respectively), as well as a muscle-specific knockdown driven by *mef2*-Gal4 (hereafter referred to as *fliI*-*KD*). Consistent with previous reports, all three conditions resulted in complete flightlessness at 24 hours after eclosion (AE) (Fig. 1b; SFig. 1a) (Campbell et al., 1993; Deak et al., 1982; Deng et al., 2021; Miklos and De Couet, 1990; Perrimon et al., 1989; Schnorrer et al., 2010). Upon closer examination, we observed the presence of large myofibrillar aggregates on the surface of the otherwise normally-sized set of dorsal longitudinal muscle (DLM) fibers. Higher-resolution imaging further disclosed that individual myofibrils are thinner than in wild type, and appear disorganized with severely disrupted sarcomeric units and an absence of regular actin and Myosin banding patterns (Fig. 1c; SFig. 1b, 2). In addition, Kettin and α-Actinin staining revealed irregularities in the size and spacing of the Z-discs, whereas the Obscurin distribution remained largely unaffected in the thin myofibrils (SFig. 2). The sarcomeric defects were quantified by using isolated individual myofibril preparations that permit precise measurements (Görög et al., 2025); a significant reduction in both sarcomere length and width was detected across all three hypomorphic conditions, with *fliI*-*KD* exhibiting the most pronounced effects (Fig. 1d; SFig. 1c). In contrast, muscle attachment sites and fiber cross-sectional area remained normal in these IFMs (SFig. 1d,e).

**Figure 2.**
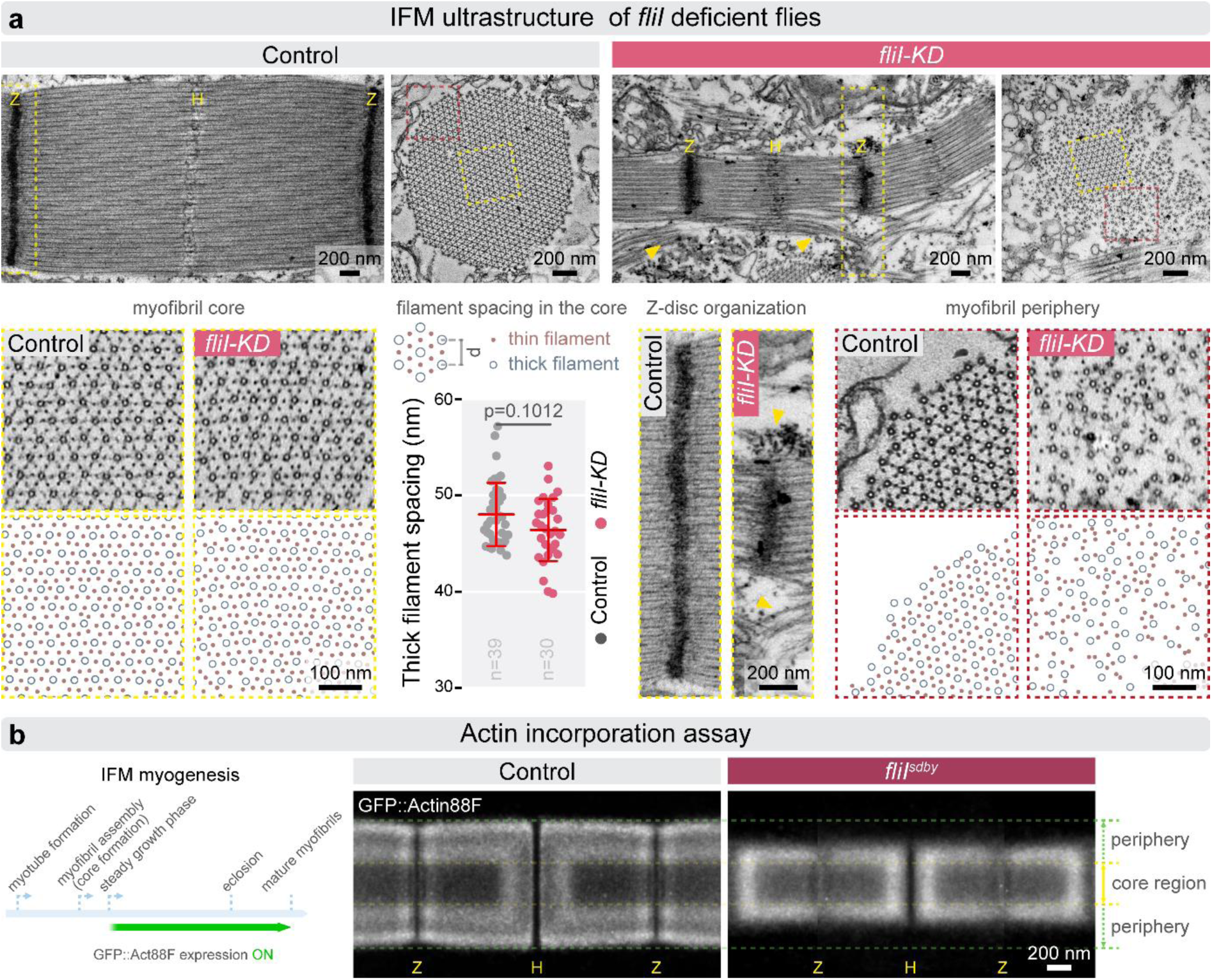
The lack of FliI affects Z-disc formation and lateral myofilament integration. **(a)** Electron micrographs of longitudinal and cross-sections of DLM fibers show that sarcomeres in *fliI-KD* (*mef2-Gal4/UAS-TRIP.JF02720*) IFM are shorter and thinner than in controls. Z-discs (yellow “Z”) frequently fail to span the full width of the myofibril and are instead restricted to the core region, leaving peripheral myofilaments partially or completely unintegrated (yellow arrowheads). Boxed regions (dotted rectangles) highlight areas enlarged below; the myofibril cores (yellow boxes; on the left) display the normal hexagonal lattice pattern, while in the peripheral areas (dark pink boxes; on the right) the myofibril cores are surrounded by unincorporated thin and thick filaments (devoid of hexagonal packing) in *fliI-KD* IFMs. Z-disc size is significantly reduced and the Z-discs often fail to reach the myofibril periphery (yellow arrowheads). Despite of these defects, Z-disc width and organization within the core appear comparable to controls. Cross sections show that the core myofilament lattice retains normal hexagonal arrangement and spacing, as indicated by unchanged average thick filament distances (p = 0.1012). Statistical significance was assessed using a Mann-Whitney test. Error bars represent mean ± s.d.; n indicates independent measurements. **(b)** Left: schematic of IFM myogenesis and experimental design of the actin incorporation assay. GFP-actin was expressed using *flightin-Gal4* (active in IFM after 48 hours APF), and incorporation into sarcomeres was assessed at ∼24 hours AE. Right: averaged dSTORM reconstructions show GFP-actin localization in control and *fliI* mutant (*fliI^stby^*) backgrounds. In controls, labeled actin incorporates at the sarcomere periphery and pointed ends, forming a frame around the core. In *fliI* mutants this peripheral incorporation is markedly reduced, whereas the unlabeled core region remains similar in size to controls. Source data are provided in Fig2SourceData.

We next examined the ultrastructure of control and *fliI* deficient IFMs (*fliI^sdby^* and *fliI-KD*) with TEM at 24 hours AE. Longitudinal and cross-sectional micrographs of DLM fibers confirmed that sarcomeres are significantly shorter and narrower in the absence of *fliI* (Fig. 2a). In addition, prominent myofibrillar aggregates were observed, frequently adopting a zebra-body-like appearance (electron-dense structures surrounded by thin filaments) (SFig. 3b), a feature also reported in *fliI*-deficient mouse hearts (Kuwabara et al., 2023), and frequently associated with various myopathies (Lake and Wilson, 1975). Interestingly, TEM also revealed that the Z-discs frequently fail to span the entire width of the myofibrils and confine to the central region of the myofibrils (Fig. 2a; SFig. 3a). This leaves a variable number of peripheral myofilaments either loosely attached or entirely disconnected from the myofibril. This phenotype is particularly apparent in cross-sectional views, which reveal a regularly organized myofibril core surrounded by unintegrated thin and thick filaments that lack the characteristic hexagonal packing (Fig. 2a; SFig. 3a). In addition, it is notable that the ends of the unintegrated peripheral myofilaments do not appear to be in register with that of the filaments stably integrated into the myofibril core (Fig. 2a; SFig. 3a). Within the core region of *fliI* mutant myofibrils, the ratio of thin to thick filaments remained at 3:1 and the spacing of the hexagonal myofilament lattice, as assessed by average thick filament distances, remained normal (Fig. 2a; SFig. 3a). These observations indicate that organization of the myofibril core is indeed largely intact, despite the core diameter being in the range of 600–900 nm, substantially below the wild-type myofibril width of approximately 1.5 µm.

**Figure 3.**
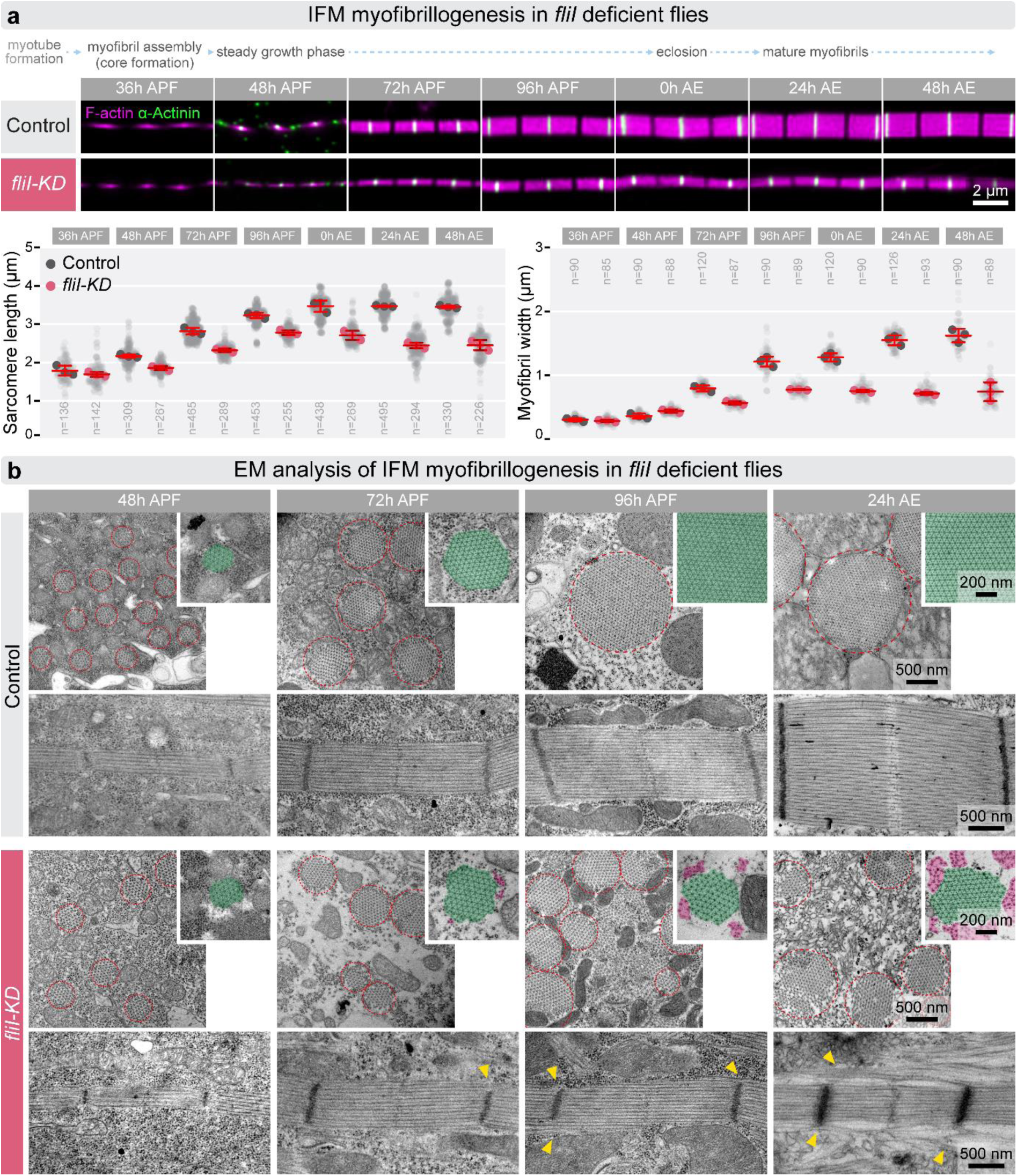
Developmental analysis of *fliI* deficient IFMs reveals a myofibril growth arrest at 60-70 hours APF. **(a)** Top: schematic outline of the timeline and key stages of IFM myogenesis. Representative images of isolated myofibrils show sarcomere growth in control and *fliI-KD* (*mef2-Gal4/UAS-TRIP.JF02720*) IFMs across seven time points (from 36 hours APF to 48 hours AE), as indicated in the schematic. F-actin (magenta) is labeled with phalloidin, and Z-discs (green) with α-Actinin. Graphs below show quantification of sarcomere length and myofibril width during development. Light gray dots represent mean values for individual measurements, and larger dots indicate means from independent experiments. The mean and s.d. of these experiments are provided, and n denotes the number of independent measurements. At 36 hours APF, no differences are observed between control and *fliI-KD*. From 48 hours APF onward, sarcomere length is significantly reduced in *fliI-KD* IFMs (p<0.0001). Myofibril width is initially comparable but becomes significantly reduced from 72 hours APF through adulthood (p<0.0001). Statistical significance was assessed using a Two-way ANOVA with Dunnett’s multiple comparison. **(b)** Electron micrographs of longitudinal and cross-sections of DLM fibers at four developmental time points show sarcomere ultrastructure in control and *fliI-KD* IFMs. Insets display high-magnification cross-sections, with regularly packed lattice regions highlighted in green and disconnected myofilaments in magenta. At 48 hours APF, mutant ultrastructure appears similar to control. By 72 hours APF, myofibrils are thinner and exhibit few disconnected myofilaments, along with emerging Z-disc defects (yellow arrowheads). At 96 hours APF, myofibrils display irregular edges and abundant disconnected filaments, resembling to the adult phenotype (24 hours AE). Raw data are provided in Fig3SourceData.

To complement the structural studies, we performed a pulse-chase experiment to directly investigate the effect of *fliI* on thin filament assembly. GFP-tagged actin monomers were expressed under the control of *fln*-Gal4, which drives expression in the IFM from 48 hours after puparium formation (APF) onward, and sarcomeric actin incorporation was evaluated at the completion of myofibrillogenesis at 24 hours AE using dSTORM nanoscopy. In wild-type animals, labeled actin was incorporated at the periphery of the sarcomeres and at the pointed end region of the thin filaments, forming the anticipated “frame” surrounding the unlabeled sarcomere core (Fig. 2b) (Gorog et al., 2025; Mardahl-Dumesnil and Fowler, 2001; Nikonova et al., 2024; Roper et al., 2005; Shwartz et al., 2016). By contrast, in the *fliI^sdby^* mutant background, this GFP::actin “frame” was markedly narrower (Fig. 2b), enclosing an unlabeled sarcomere core identical in size to that of the corresponding wild-type control (Fig. 2b).

Collectively, these findings further confirmed that FliI is essential for muscle development in fruit flies. The data indicate that loss of FliI does not prevent myofibril formation *per se*, nor does it appear to be required during the initial phases of flight muscle development, as evidenced by the normal number of muscle fibers and properly formed attachment sites (SFig. 1d,e). However, FliI is critically required during the growth phase to ensure that sarcomeres attain their mature dimensions. Moreover, we demonstrated that FliI deficiency results in the formation of numerous actin-based protein aggregates, a hallmark of various myopathies also observed in *fliI* mutant mice. Importantly, we further revealed that Z-discs are frequently confined to the central myofibril region in *fliI* mutants, a defect that is accompanied by a failure of stable peripheral myofilament integration. Taken together, these results suggest that the primary contribution of FliI to myofibril growth is not the promotion of myofilament formation, instead it emerges as a specific factor required for the integration of myofilaments at the Z-disc.

### FliI is required for myofibril maturation

To analyze the developmental role of FliI in greater details, we examined isolated individual myofibrils from seven defined stages of myofibrillogenesis by confocal microscopy. At our standard laboratory conditions at 25°C the first immature myofibrils (or premyofibrils) assemble by 32-36 hours APF, and accordingly 36 hours APF was selected as the earliest developmental time point, followed by three additional pupal (48, 72 and 96 hours APF) and three adult (0, 24 and 48 hours AE) time points. Following their initial assembly at 36 hours APF, myofibrils contain about 100 sarcomeres; by 48 hours APF this number increases to approximately 230 sarcomeres (Rodier et al., 2025; Spletter et al., 2018). Thereafter, only a modest number of sarcomeres are added, reaching about 270 sarcomeres per myofibril at 60 hours APF, a value that remains stable until eclosion (Spletter et al., 2018). Beyond the changes in sarcomere number, myofibrils undergo a substantial growth that proceeds in two major phases. During the first phase (36-48 hours APF), premyofibrils adopt a more organized structure, establishing the lattice spacing characteristic of mature myofibrils, with thin and thick filaments arranged in the hexagonal array typical of adult muscle (Gorog et al., 2025); however, no major changes in sarcomere dimensions occur during this period (Gorog et al., 2025; Orfanos et al., 2015; Reedy and Beall, 1993; Spletter et al., 2018), we therefore designate this interval the “organization phase.” The second phase, spanning 48 hours APF to 24 hours AE, is characterized by a continuous increase in sarcomere length and diameter until mature dimensions are attained (Gorog et al., 2025; Orfanos et al., 2015; Reedy and Beall, 1993; Spletter et al., 2018); and therefore we refer to this interval as the “steady growth phase”. Myofibrils isolated from 36 hours APF showed no significant differences between control and *fliI* mutant IFMs with respect to sarcomere length or width (Fig. 3a; SFig. 4a). From 72 hours APF onward through the adult stages, however, *fliI*-deficient sarcomeres were significantly shorter than controls. While at 48 hours APF we found no significant differences in sarcomere width, the lack of FliI caused a decrease in myofibril width from 72 hours APF till adulthood (Fig. 3a; SFig. 4a). A similar developmental analysis has been carried out with TEM to collect higher resolution data and to examine entire muscle fibers. At 48 hours APF, the ultrastructure of *fliI* mutant muscle fibers was indistinguishable from controls (Fig. 3b). By 72 hours APF, however, thinner myofibrils were clearly evident, and a small number of disconnected myofilaments could be detected in both longitudinal and cross sections (Fig. 3b), accompanied by the characteristic Z-disc defects previously described in adult muscles (Fig. 3b). By 96 hours APF, the myofibrils exhibited the same abnormal morphology with irregular edges as observed in adults, and disconnected myofilaments were present in substantial amounts near all myofibrils (Fig. 3b), similar to the phenotype observed at 24 hours AE. This temporal progression clearly indicates that these defects arise from developmental abnormalities rather than from muscle disorganization caused by contractile activity. Furthermore, quantitative comparison of thick filament numbers demonstrated that the diameter of the myofibril core (defined by regions exhibiting normal hexagonal packing) remains approximately constant from 72 hours APF through 24 hours AE in *fliI*-*KD* animals (Fig. 3b), indicating that peripheral filament integration is arrested at or before 72 hours APF.

**Figure 4.**
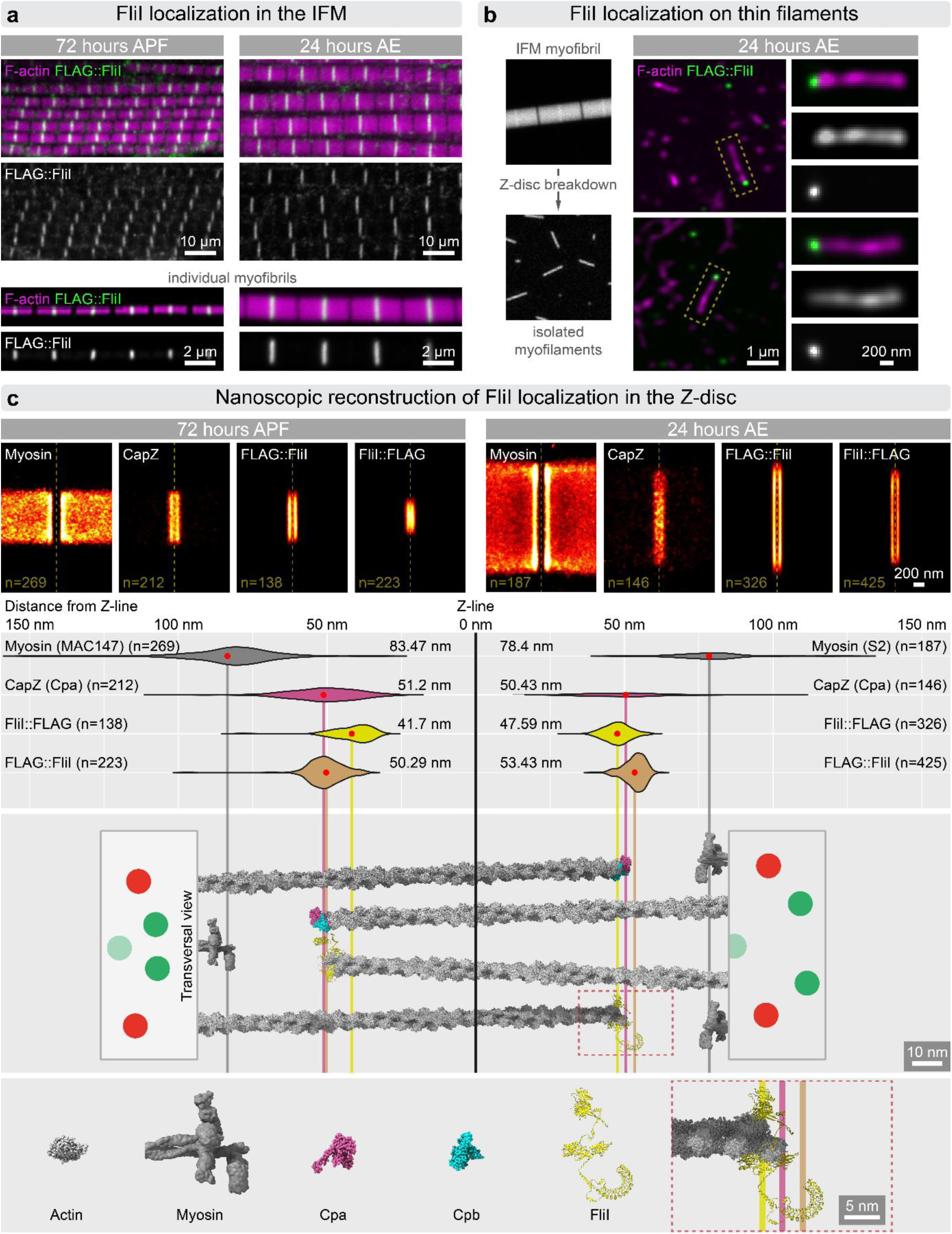
The FliI protein accumulates at the barbed end of the actin filaments, located at the edges of the Z-discs. (**a**) Localization of FLAG::FliI in IFMs of *fliI-null* rescue animals, shown in thoracic sections (top) and isolated myofibrils (bottom) at developing (left) and mature (right) stages. F-actin (magenta) is labeled with phalloidin, and FLAG::FliI is detected via FLAG staining (green or gray), localizing exclusively to Z-discs in all conditions. (**b**) Left: schematic illustrating the isolation of individual thin filaments from IFM myofibrils. Right: micrographs of fragmented thin filaments from *fliI-null* rescue animals. Insets highlight thin filaments with discrete FLAG signal at their ends. (**c**) Top: Averaged dSTORM reconstructions from pupal (left) and adult (right) IFMs display the localization of C- and N-terminally FLAG-tagged FliI, alongside Myosin (MAC147) and CapZ (Cpa), which serve as reference markers for the I-band and Z-disc edges. The Z-line is indicated by a dotted yellow line; n denotes the number of sarcomeres used for averaging. Bottom: longitudinal epitope distributions relative to the Z-line, shown as violin plots with mean values indicated (red dots). A scaled molecular model of the Z-disc is aligned to the plots, with vertical lines marking average epitope positions. FliI localizes to a position consistent with capping the barbed ends of thin filaments in both pupal and adult sarcomeres. Schematic cross-sections (left and right) illustrate the thin filament lattice (red and green). The structural model of FliI was generated using AlphaFold. Visualization was performed in ChimeraX, incorporating the 7PDZ PDB structure, as well as the EMD-3301 and EMD-8727 density maps. Source data are provided in Fig4SourceData

This developmental analysis confirmed that the depletion of *fliI* does not significantly impair myofibril growth until the end of the organization phase at 48 hours APF. During the subsequent steady growth phase, however, both sarcomere elongation and the lateral thickening of myofibrils are clearly compromised. The main sarcomeric parameters appear to remain essentially constant from 72 hours APF through 24 hours AE, retaining values characteristic of the 72-hour APF stage, indicating that myofibril growth, and in particular lateral addition of new filaments, is severely affected after this point. The first loosely connected peripheral myofilaments appear between 48 and 72 hours APF, and their number increases progressively through 24 hours AE. This pattern demonstrates that the formation of myofibrillar components is not prevented in the absence of FliI; rather, their integration into the myofibrillar lattice is specifically impaired. These observations therefore corroborate that FliI is a key factor in anchoring and organizing peripheral myofilaments in the *Drosophila* IFM.

### FliI exhibits a Z-disc accumulation at the barbed end of the thin filaments

To determine the subcellular distribution of FliI in the IFM, we employed a rescue-based localization strategy using the *fliI^CRIMIC.TG4^* allele, predicted to be a protein null (Lee et al., 2018). In this system, the CRIMIC.TG4 cassette drives GAL4 expression under the control of the endogenous *fliI* promoter, thereby enabling expression of rescue constructs (Lee et al., 2018). Hemizygous males carrying this allele die during the larval (52%) or early pupal stage (48%) (SFig. 5a). When a full-length *UAS-fliI* transgene bearing either an N- or C-terminal FLAG tag was expressed in this background, the lethal phenotype was rescued till adulthood in approximately 77% of offspring (SFig. 5a). Flight ability and flight muscle organization were fully restored in these animals (SFig. 5b), demonstrating that neither tag interferes with protein function and validating this system for localization studies. Anti-FLAG immunostaining on sagittal sections of pupal (72 hours APF) and adult (24 hours AE) thoraces revealed that FliI localizes exclusively to the Z-discs of DLM myofibrils, as demonstrated by the staining pattern of the N-terminally tagged construct (Fig. 4a). These results are consistent with the heterologous antibody staining reported by Deng et al. (2021), and were further confirmed in isolated myofibrils from both adult (24 hours AE) and pupal (72 hours APF) muscles (Fig. 4a). To resolve the sarcomeric localization of FliI at higher resolution, we applied dSTORM nanoscopy combined with structure averaging (Szikora et al., 2020a). We found that FliI exhibits a distinctive “double-line” pattern along the Z-line in myofibrils from both pupal (72 hours APF) and adult (24 hours AE) IFMs (Fig. 4c). The mean Z-line peak-to-peak distances were 50.29 nm and 41.7 nm in developing IFMs (72 hours APF), and 53.43 nm (N-terminal FLAG) and 47.59 nm (C-terminal FLAG) in mature IFMs (24 hours AE). These values closely correspond to those measured for Cpa (Capping protein α) (51.20m; 50.43nm) (Fig. 4c), a marker of actin filament barbed ends and thus of Z-disc edges. Anti-Myosin heavy chain (Mhc) staining served as a reference for I-band positioning, yielding Z-line peak distances of 83.47 nm and 78.4 nm in adult and pupal myofibrils, respectively (Fig. 4c). To further assess whether FliI is genuinely associated with thin filament barbed ends in the IFM, we isolated individual myofilaments following enzymatic digestion of Z-discs (Fig. 4b). In these preparations, thin filaments were largely fragmented, yet FliI was consistently detected at filament ends. Importantly, FliI was never observed along filament sides or at both ends simultaneously (Fig. 4b), demonstrating that its Z-disc-restricted localization is not an artifact of limited epitope accessibility in the densely packed IFM.

**Figure 5.**
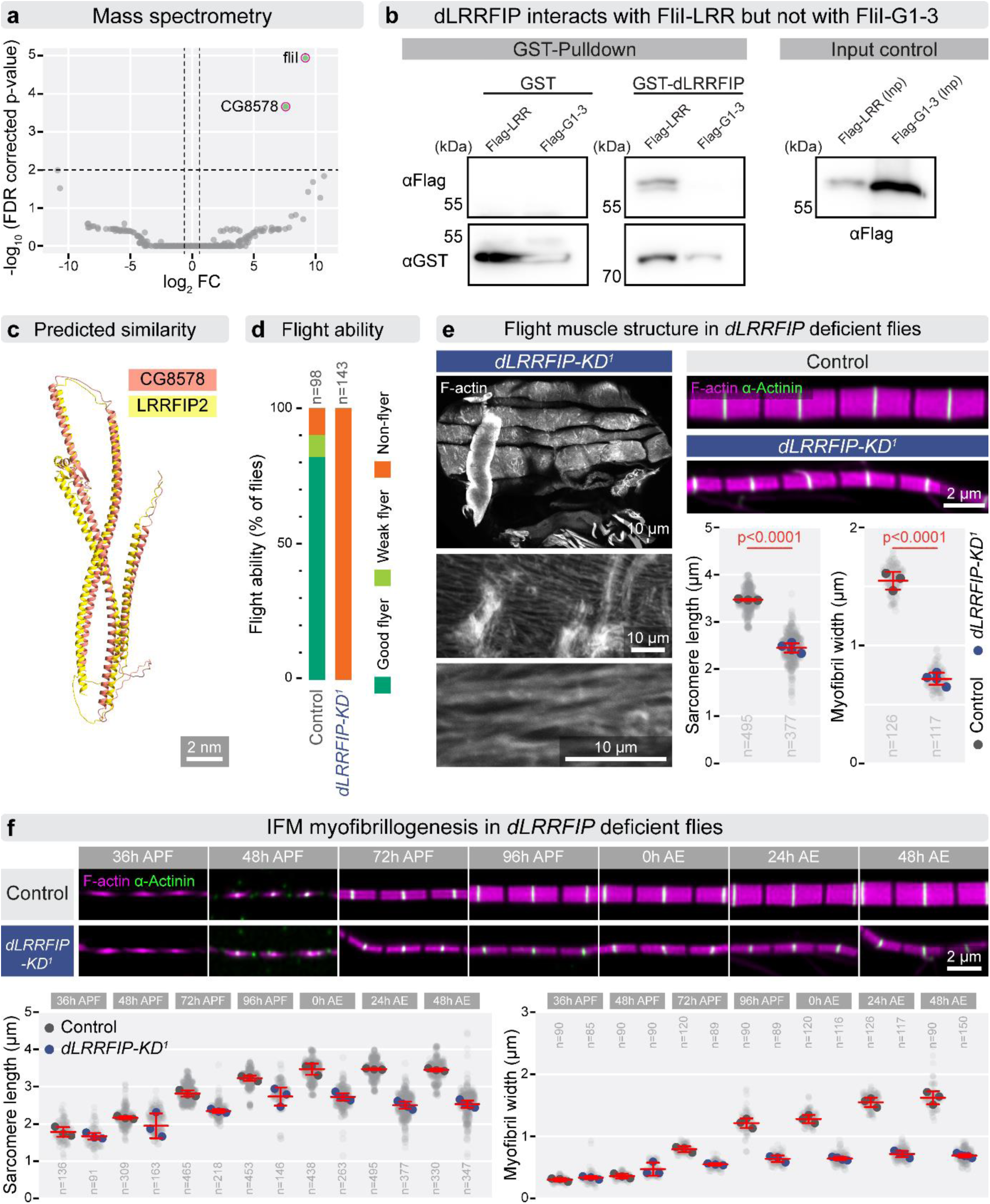
FliI interacts with dLRRFIP, the lack of which results in nearly identical IFM phenotypes as that of *fliI-KD*. (**a**) Volcano plot shows proteins enriched (green dots) by FliI-FLAG immunaffinity purification from thoracic tissue, identified by mass spectrometry. (**b**) GST pull-down assay demonstrating that GST-dLRRFIP pulls down FLAG-tagged FliI-LRR, but not FliI-GH1-3. (**c**) Image shows the structural similarity between the AlphaFold predicted structure of CG8578 and human LRRFIP2. (**d**) Flight assay showing complete loss of flight in dLRRFIP*-KD^1^* (*mef2-Gal4/UAS-KK101150*) adult flies at 24 hours AE, compared with ∼90% flight ability in controls (*w^1118^*). (**e**) On the left low- and high-magnification images of thoracic sections reveal DLM fiber structure in *dLRRFIP-KD^1^* animals, large myofibrillar aggregates are visible on the surface of otherwise normally sized fibers. At higher resolution, individual myofibrils appear thinner, disorganized, and exhibit severely disrupted sarcomeric structure with loss of the regular actin pattern. F-actin is visualized by phalloidin staining (gray). On the right, panels show isolated myofibrils from control and *dLRRFIP-KD^1^* IFMs, demonstrating reduced sarcomere length and myofibril diameter in knockdown animals. F-actin (magenta) is labeled with phalloidin, and Z-discs (green) with α-Actinin. The graph below presents morphometric quantification, confirming significant reductions in both parameters (p<0.0001). Statistical significance was assessed using unpaired t-tests. (**f**) Representative images of isolated myofibrils show sarcomere growth in control and *dLRRFIP-KD^1^* IFMs across seven time points (from 36 hours APF to 48 hours AE). F-actin (magenta) is labeled with phalloidin, and Z-discs (green) with α-Actinin. Graphs below show quantification of sarcomere length and myofibril width during development. Light gray dots represent mean values for individual measurements, and larger dots indicate means from independent experiments. The mean and s.d. of these experiments are provided, and n denotes the number of independent measurements. At 36 and 48 hours APF, no differences are observed between control and *dLRRFIP-KD^1^*. From 72 hours APF onward, sarcomere length is significantly reduced in *dLRRFIP-KD^1^* IFMs (p<0.0001). Myofibril width is initially comparable but becomes significantly reduced from 72 hours APF through adulthood (p<0.0001). Statistical significance was assessed using a Mixed-effects Model analysis with Sidak’s multiple comparison. Raw data are provided in Fig5SourceData.

Thus, these localization studies establish FliI as a Z-disc-associated protein with a pronounced accumulation at the barbed ends of actin filaments. This finding is fully consistent with our previous *in vitro* demonstration of barbed-end capping activity by the gelsolin homology domains of *Drosophila* FliI (Pintér et al., 2020). Furthermore, the Z-disc enrichment is also in harmony with the Z-disc phenotypes observed upon *fliI* loss, and collectively these data support a model in which FliI functions as a barbed-end-associated factor crucial for peripheral myofilament integration at the Z-disc.

### FliI interacts with dLLRFIP in the flight muscle of *Drosophila*

To gain further insight into the molecular mechanisms of FliI, we aimed to identify its interaction partners in the IFM. Human FliI has formerly been shown to interact with numerous proteins across diverse tissues and cell types (for a comprehensive review, see Strudwick and Cowin, 2020). To identify *Drosophila* IFM-specific interactors, we performed affinity purification coupled to mass spectrometry using thoracic protein extracts from flies expressing FLAG-tagged FliI in a protein-null background (*fliI^CRIMIC.TG4^)*. Restricting the analysis to thoracic extracts enriched for IFM-specific interactions, and following two biological replicates, a single strong candidate interactor was identified, the product of the *CG8578* gene (Fig. 5a). Although *CG8578* remains largely uncharacterized in *Drosophila*, an interaction between FliI and this gene product has previously been reported in *Drosophila* S2R+ cells (Guruharsha et al., 2011), and its mammalian orthologs are likewise known to interact with FliI (Fong and de Couet, 1999; Liu and Yin, 1998). The two human homologs of *CG8578*, LRRFIP1 and LRRFIP2, were originally isolated from HeLa cells as FliI leucine-rich repeat (LRR) domain-interacting proteins (Fong and de Couet, 1999). As the *Drosophila* genome appears to encode a single gene with significant homology to both human paralogs, *CG8578* will hereafter be referred to as *Drosophila* LRRFIP (dLRRFIP; Fig. 5c). The LRR domain-specific interaction between *Drosophila* FliI and dLRRFIP was independently confirmed by a GST pull-down assay (Fig. 5b), supporting a highly conserved mode of interaction.

We next asked whether dLRRFIP is required during muscle development, and we found that muscle-specific knockdown of dLRRFIP (using two independent *UAS-IR* lines, referred to as KD^1^ and KD^2^) resulted in complete loss of flight ability and severe myofibril disorganization (Fig. 5d,e; SFig. 6b,c). Knockdown of *dLRRFIP* led to the formation of large actin aggregates (Fig. 5e), accompanied by reductions in sarcomere length and myofibril width (Fig. 5e; SFig. 6c), closely mirroring the phenotype observed upon *fliI*-*KD* at 24 hours AE. Developmental analysis of *dLRRFIP* silencing yielded similarly concordant results: loss of dLRRFIP had no significant effect on myofibril growth through 48 hours APF, at which point sarcomere length and width were indistinguishable from wild type; however, both parameters were significantly reduced in the KD^1^ line by 72 hours APF (Fig. 5f). This reduction persisted through all subsequent stages examined (up to 48 hours AE) (Fig. 5f), and as observed for *fliI*, loss of dLRRFIP appears to arrest significant growth in both length and width after 72 hours APF. As expected, TEM analysis of the *dLRRFIP-KD^1^* muscles revealed nearly identical phenotypic effects as for *fliI*, including the presence of actin aggregates (frequently adopting zebra body-like electron-dense structures) (SFig. 6a), Z-disc malformations characterized by centrally confined narrow discs, and a high abundance of unintegrated peripheral myofilaments (SFig. 6a). Collectively, the biochemical evidence and the striking similarity of phenotypic outcomes provide strong support for FliI and dLRRFIP acting together during muscle development.

**Figure 6.**
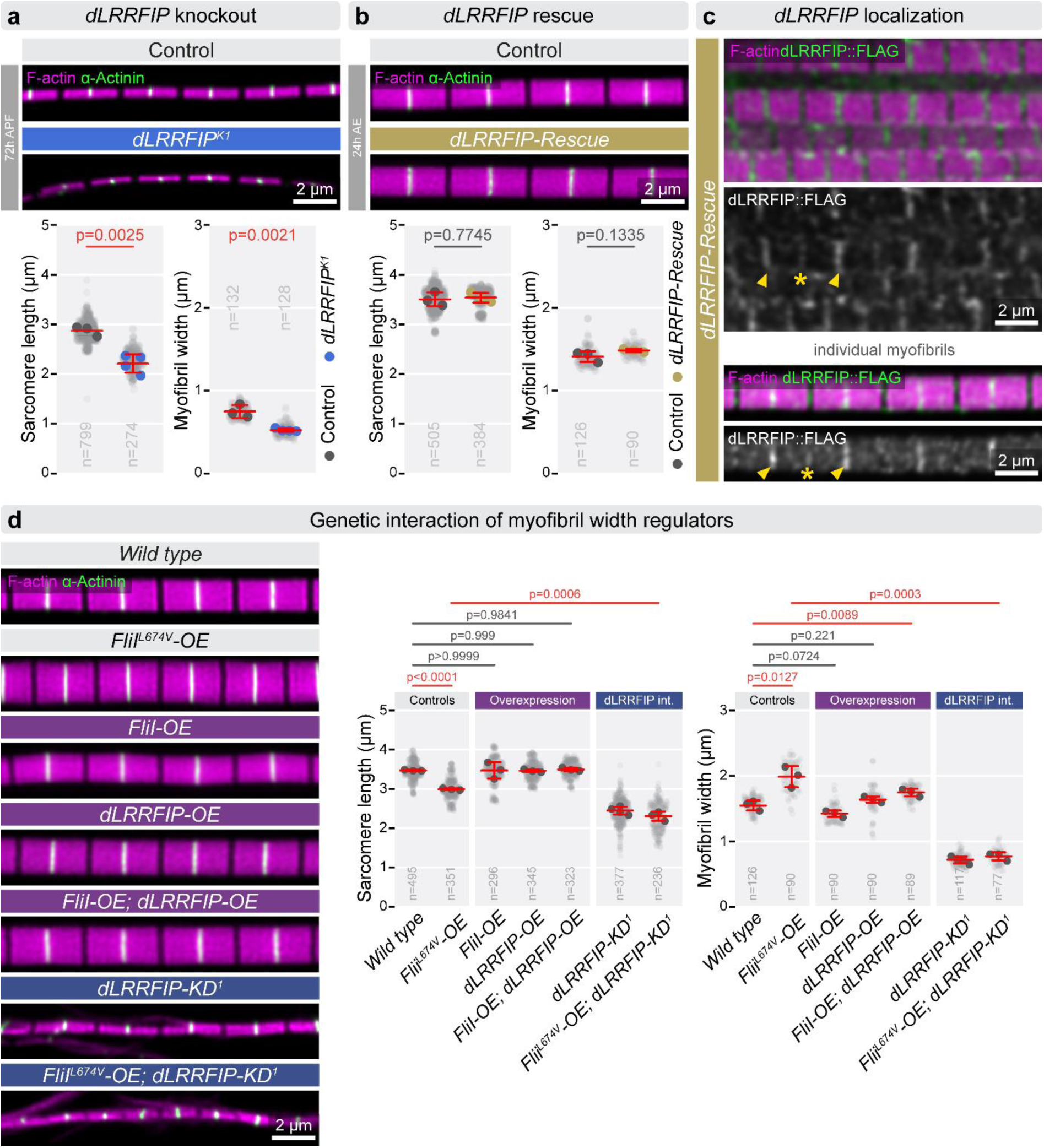
dLRRFIP localization in the IFM, and genetic epistasis analysis with FliI^L674V^. **(a)** Analysis of individual myofibrils from *dLRRFIP^K1^* null mutant pupal IFMs (72 hours APF) reveals a reduction in sarcomere length (p=0.0025) and width (0.0021), that can be rescued by muscle specific expression of dLRRFIP::FLAG (p=0.7745 and 0.1335, respectively) (**b**). Statistical significance was assessed using unpaired t-tests. F-actin (magenta) is labeled with phalloidin, Z-discs (green) are marked with α-Actinin. (**c**) The localization pattern of dLRRFIP::FLAG (in green) in rescue conditions in hemi-thoraces (upper panels) and in individual myofibrils (lower panels), displaying an accumulation at the Z-discs (arrowheads) and in the H-zone (asterisk). (**d**) (*left*) Individual myofibrils stained for actin (magenta) and α-Actinin (green) to mark the Z-discs from adult 24 hours AE IFMs of the genotype indicated. (*right*) Whereas, overexpression (OE) of wild type FliI does not result in a significant increasement in sarcomere width (and length), the OE of dLRRFIP or the concomitant OE of FliI and dLRRFIP induces a moderate increase in myofibril width (without a notable alteration in length). By contrast, FliI^L674V^ strongly increases myofibril width, producing a GOF type effect in this regard, which is completely suppressed by *dLRRFIP-KD^1^*. Light gray dots represent individual measurements, while larger dots indicate mean values from independent experiments. Error bars show mean of independent experiments ± s.d., and n denotes the number of independent measurements. Statistical significance was assessed either using unpaired t-tests or One-way ANOVA with Dunnett’s multiple comparison. Raw data are provided in Fig6SourceData.

The existence of a FliI-dLRRFIP complex implies that these proteins share overlapping expression patterns in the IFM. To test this, we generated a *dLRRFIP* null allele (*dLRRFIP^K1^*) by CRISPR/Cas9-mediated deletion of the entire coding sequence, which was subsequently combined with a *UAS-dLRRFIP-FLAG* transgene in a rescue configuration. The *dLRRFIP^K1^* null allele is pupal lethal, enabling structural analysis of the IFM at 72 hours APF, where it produced phenotypic effects identical to those observed upon muscle-specific knockdown (Fig. 6a, SFig. 7a). Ubiquitous expression of *UAS-dLRRFIP-FLAG* driven by *act5C*-Gal4 in this mutant background was sufficient to partially rescue lethality and to fully restore flight ability (in 19 of 21 individuals) and myofibril organization (Fig. 6b). Determination of the dLRRFIP protein distribution pattern in this rescue context revealed accumulation at both the Z-disc and the H-zone (Fig. 6c). Thus, FliI and dLRRFIP are both present at the Z-disc, and their genetic impairment in multiple ways consistently produces highly similar, if not identical, defects in IFM development.

**Figure 7.**
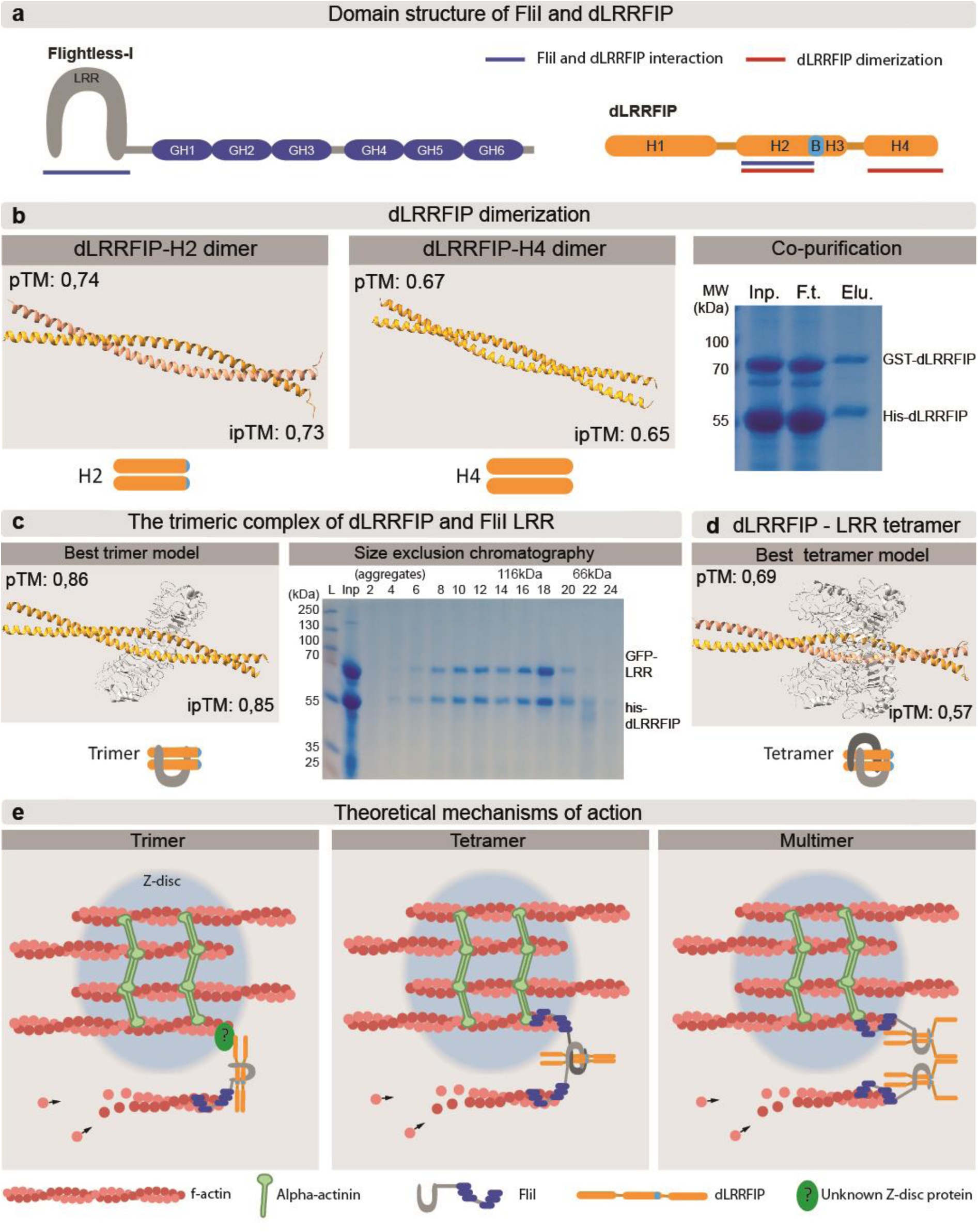
FliI and dLRRFIP form a complex. (**a**) Domain structure of the FliI and dLRRFIP proteins, the LRR and gelsolin homology (GH1-GH6) domains of FliI, as well as the helix domains (H1-H4) of dLRRFIP are indicated. Regions predicted to be involved in intermolecular interactions are highlighted. (**b**) Molecular models show that the H2 and H4 helices of dLRRFIP are able to dimerize as predicted by AlphaFold2, and accordingly, GST tagged dLRRFIP co-purifies in a 1:1 ratio with its Histidine tagged counterpart during metal affinity purification, indicating the formation of a stable homo dimer. Inp.: input, F.t.: flow through, Elu.: elution. (**c**) Based on *in silico* modelling, the LRR domain of FliI LRR binds the dimeric H2 helix of dLRRFIP. GFP-LRR and dLRRFIP are co-purified during size exclusion chromatography indicating the existence of a stable complex. Proteins in fraction 12 correspond to the expected size of a trimeric complex (169 kDa). Note the presence of larger complexes in fractions 6-10, near the exclusion volume, L: protein weight ladder, Inp: input. (**d**) Molecular model of two dLRRFIP H2 helices and two FliI-LRR forming a tetrameric structure with reasonable high pTM and ipTM scores. (**e**) Based on *in silico* and biochemical data, we envision three possibilities whereby the FliI-dLRRFIP complex could incorporate a new actin filament at the Z-disc. In each scenario the FliI GH domains are responsible for binding of an actin filament, while the LRR domain binds a dimeric dLRRFIP. In the first case a trimeric FliI-dLRRFIP complex localizes to the Z-disc through an unknown Z-disc protein while binding to an actin filament that ensures the lateral growth. In the second case, a tetrameric complex binds an already incorporated actin filament that directs the complex to the Z-disc while also stabilizing a new filament through the GH domains of the second FliI protein of the complex. In the third scenario multiple FliI-dLRRFIP trimers could form a larger complex, capable of binding already incorporated and newly recruited/laterally integrated actin filaments, similar to the tetrameric structure in the former case.

### dLRRFIP is essential for the proper FliI function

To address the functional relationship between FliI, dLRRFIP, and previously identified regulators of lateral sarcomere growth, we employed a series of genetic epistasis experiments. Overexpression of either FliI alone in a wild-type background had no significant effect on myofibril width (Fig. 6d), a result consistent with FliI being subject to autoinhibition (Nag et al., 2013), whereas the excess of dLRRFIP alone or the concomitant overexpression of both FliI and dLRRFIP produced a moderately strong myofibril-widening effect, although only the latter appears to be statistically significant (Fig. 6d). By contrast, expression of the FliIL674V point mutant form (with an impairment in the GH2 domain) (Ruijmbeek et al., 2023) elicited a strong gain-of-function (GOF) effect on myofibril width (Fig. 6d), providing a sensitized background well suited for genetic epistasis analysis. The GOF effect of FliIL674V was completely suppressed by knockdown of dLRRFIP (Fig. 6d), demonstrating that FliI activity is dependent on the presence of its major interaction partner. The myofibril widening phenotype of FliIL674V was also entirely suppressed by loss of the formin Fhos, and knockdown of DAAM, a second formin, produced a strong suppression as well (SFig. 7b). Although the KD of Zasp52 had a negligible effect on FliIL674V (SFig. 7b), and the KD of SALS in this background resulted in larval lethality, precluding its assessment, these observations further substantiate the involvement of the previously identified components of lateral myofibril growth, and collectively the data are consistent with a reiterative mechanism in which all components are required for the sequential addition of new peripheral myofilament layers.

### Molecular basis of the FliI/LRRFIP interaction

To gain insight into the molecular basis of the FliI/dLRRFIP interaction and its potential biological significance, we combined biochemical assays using heterologously expressed proteins with a structural modelling approach employing AlphaPulldown (Molodenskiy et al., 2025; Yu et al., 2023). Previous studies on human LRRFIP1 have established that it forms a homodimer via its coiled-coil region (Nguyen and Modis, 2013), which is structurally analogous to Helix2 of dLRRFIP. Consistent with this, structural predictions indicate that Helix2 and Helix4 of dLRRFIP likewise mediate dimer formation (Fig 7a,b). This prediction is supported by our experimental data as dLRRFIP proteins with different affinity tags were co-purified in a 1:1 ratio, and dimer formation was observed during gel filtration as well (Fig. 7b, SFig. 8b).

**Figure 8.**
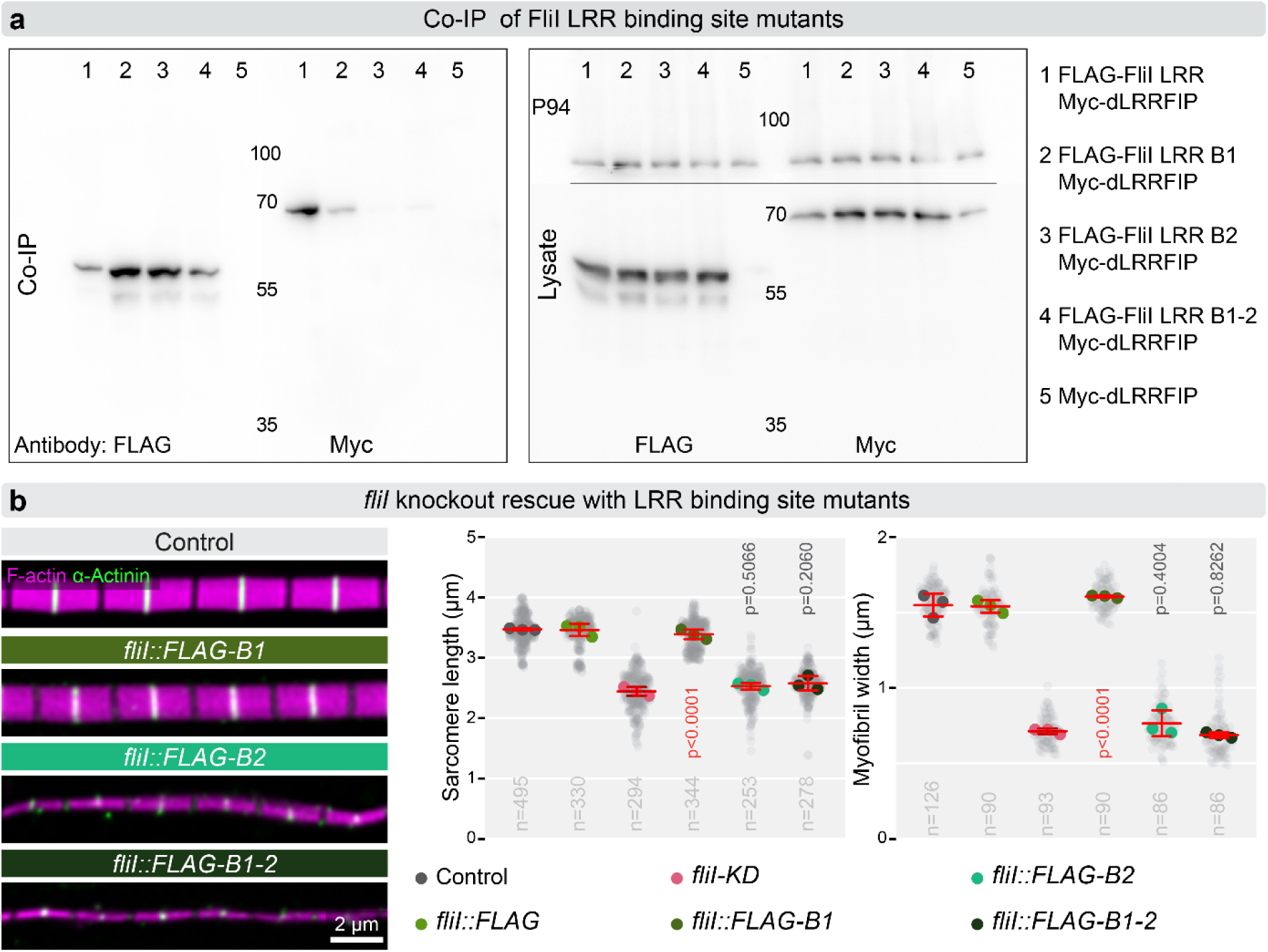
Mutations affecting the binding of FliI LRR to dLRRFIP impair muscle development. (**a**) (Left panel) A co-immunoprecipitation experiment with S2 cell lysates reveals that the B1 mutant version of FliI LRR exhibits a reduced dLRRFIP binding, while the B2 and B1-2 mutations completely abolish that. The FLAG antibody was able to bind the LRR variants (Co-IP, FLAG). (Right panel) All proteins were expressed in comparable levels, and were detectable in the cell lysates (Loading control: P94). (**b**) Rescue of the *fliI* null allele with control and dLRRFIP binding site mutants. Confocal imaging (on the left), followed by sarcomere size measurements (on the right), revealed that the B1 mutant is able to restore wild type muscle structure, while the B2 and B1-2 mutants fail to rescue. Light gray dots represent individual measurements, while larger dots indicate mean values from independent experiments. Error bars show mean of independent experiments ± s.d., and n denotes the number of independent measurements. Statistical significance was assessed using One-way ANOVA with Dunnett’s multiple comparison. Raw data are provided in Fig8SourceData.

Having established that dLRRFIP dimerizes, we turned to characterizing the interaction between the LRR domain of FliI and dLRRFIP. Gel filtration assays demonstrated that bacterially expressed eGFP-tagged FliI LRR domain co-elutes with dLRRFIP, forming at minimum a tripartite complex; and higher-order assemblies were also detected (Fig. 7c; SFig. 8b,e–g). Structural modelling of the putative trimeric complex, comprising the FliI LRR domain and a dLRRFIP dimer, consistently yielded high-confidence models in which dLRRFIP dimerizes via Helix2, with the resulting interface engaging the concave inner surface of the FliI LRR domain (Fig. 7c; SFig. 8b). To identify the key residues mediating the FliI LRR–dLRRFIP interaction, we applied ChimeraX contact-finding algorithms in conjunction with PDBePISA (Krissinel and Henrick, 2007), which revealed two discrete interaction surfaces on the LRR domain, designated B1 and B2 (SFig. 9). Alanine substitution of the critical residues at B1 (B1Ala) partially reduced the FliILRR-dLRRFIP interaction, whereas the B2Ala and B1-2Ala double mutants completely abolished it, as assessed by co-immunoprecipitation in S2 cells (Fig. 8a). Consistent with these biochemical findings, the B2Ala and B1-2Ala mutant forms of FliI failed to rescue the *fliI^CRIMIC.TG4^* null allele (Fig. 8b), providing strong *in vivo* validation of the structural predictions.

During modelling of the FliI LRR–dLRRFIP interaction, we identified the possibility of a higher-order double-dimeric assembly composed of two FliI LRR domains bound to a single dLRRFIP dimer (Fig. 7d, SFig. 8c). Although the confidence score for this molecular model is lower than for the trimeric configuration, we note that such a FliI–dLRRFIP double dimer would generate a complex presenting two actin barbed-end binding surfaces, contributed by the GH domains of each FliI subunit. Combining the *in silico* predictions with the experimental data, we conclude that formation of the FliI–dLRRFIP complex is absolutely required for peripheral thin filament integration. We propose that the LRR domain mediated interaction of FliI and dLRRFIP requires prior dimerization of dLRRFIP, and that the resulting complex either functions as a structural mechanical link in its trimeric form (Fig. 7e), or assembles into higher-order complexes presenting multiple actin-binding sites capable of crosslinking thin filament barbed ends (Fig. 7e).

## DISCUSSION

Recent research has identified *fliI* as a genetic factor in cardiomyopathy (Al-Hassnan et al., 2020; Karczewski et al., 2020; Kuwabara et al., 2023; Lipov et al., 2023; Ruijmbeek et al., 2023). While previous studies have provided some insights into the muscle-specific function of FliI, a comprehensive molecular model explaining its mode of action is still lacking. In this study, we used a multifaceted approach to uncover the molecular mechanism by which the conserved FliI protein regulates myofibrillar structure and growth.

Although the founding member of the FliI protein family was originally identified in *Drosophila* via mutations that impair flight ability and myofibrillar organization of the IFM (Campbell et al., 1993; Perrimon et al., 1989), it took until lately to demonstrate that FliI plays a widespread role in muscle development and it is required for heart morphogenesis in zebrafish, mouse and human (Kuwabara et al., 2023; Ruijmbeek et al., 2023). By analyzing a limited number of DCM (dilated cardiomyopathy) patient derived alleles alongside with *de novo* generated loss of function alleles, these studies reached at least partly conflicting conclusions regarding the role of FliI. Notably, Kuwabara and colleagues suggested that FliI regulates sarcomeric actin dynamics, and based on postnatal conditional knockout experiments, reported a moderate reduction in sarcomere length (Kuwabara et al., 2023). In contrast, Ruijmbeek and colleagues reported a complex heart phenotype with severe structural defects in cardiomyocytes, affecting cell adhesion and myofibril organization, and heart trabeculation (Ruijmbeek et al., 2023), and proposed that the primary role of FliI may be in linking myofibrils to the plasma membrane through Vinculin and N-cadherin adhesion complexes. Furthermore, a recent *Drosophila* study connected the IFM myofibrillar defects caused by *fliI* knockdown to an actin severing activity (Deng et al., 2021). In contrast to all these models, we show here that FliI is primarily required for lateral integration of myofilaments during myofibril maturation in the *Drosophila* flight muscle. Although Vinculin is known to form distinct foci at myofibril attachment sites adjacent to epidermal tendon cells in the IFM (Green et al., 2018; Maartens et al., 2016), myofibril attachment sites remain normal in *fliI*-deficient animals, indicating that *Drosophila* FliI does not play a role in myofibril attachment. Regarding the severing model, Gelsolin proteins were indeed shown to sever actin filaments (Barrie et al., 2025; Burtnick et al., 1997; Chaponnier et al., 1986), and a severing activity has been proposed for C. elegans FliI based on TEM observations (Goshima et al., 1999). However, actin severing activity could not be confirmed for mouse FliI in biochemical experiments (Mohammad et al., 2012), and the GH domains of *Drosophila* FliI similarly lack this activity (Pintér et al., 2020). Instead, barbed-end capping has been identified as the principal biochemical activity of FliI proteins (Arora et al., 2015; Mohammad et al., 2012), clearly distinguishing them from canonical Gelsolin superfamily members. Sarcomeres in *fliI* knockdown IFM myofibrils are shorter than in wild type, which parallels observations in mouse cardiac myofibrils and was interpreted as evidence for a role in thin filament elongation (Kuwabara et al., 2023). However, this interpretation is difficult to reconcile with biochemical data demonstrating that FliI inhibits actin assembly *in vitro* (Arora et al., 2015; Mohammad et al., 2012; Pintér et al., 2020). While we cannot exclude that mouse FliI exhibits a different activity *in vivo* in the context of its protein interaction partners, we consider it most likely that the concomitant reduction in both sarcomere width and length in the *Drosophila* IFM reflects a coordination between lateral and longitudinal sarcomere growth. Given that the pointed end region (H-zone) is thought to be the principle site of thin filament elongation (Littlefield et al., 2001; Szikora et al., 2022), yet FliI accumulates at barbed ends within the Z-disc, we propose that FliI is primarily involved in peripheral myofilament addition (increasing myofibril width), and that the reduction in sarcomere length observed upon loss of FliI is an indirect consequence.

Several decades of research across model organisms and myocyte cultures have established that the fundamental mechanisms of myofibrillogenesis are highly conserved throughout the animal kingdom. Although numerous alternative models have been considered, the three-step premyofibril model has become broadly accepted as an explanation for how muscle-specific proteins assemble into series of functional sarcomeres (Sanger et al., 2017). While the precise sequence of events may vary across the wide range of muscle types, a common theme is that the premyofibrillar, premature sarcomeric units undergo a substantial growth through lateral addition of myofilaments to increase myofibril diameter. Despite the importance of this process, how new myofilaments form and how they become integrated into the developing myofibrillar lattice have remained largely unexplored. Here, we show that in the absence of FliI or its interaction partner dLRRFIP, morphologically normal myofibril cores form, but as development proceeds, a growing number of myofilaments accumulate at the periphery of the myofibril cores and fail to adopt a regular arrangement or integrate into the core region. This phenotype directly implicates the FliI/dLRRFIP complex in the lateral incorporation of nascent myofilaments. Since FliI is an actin binding protein, we propose that it plays a direct role in thin filament integration, which is subsequently followed by myosin/thick filament integration. In the IFM, FliI is exclusively enriched at the edges of the Z-disc, and consistent with a concerted action, dLRRFIP is also present at the Z-disc — highlighting the Z-disc as a key site of peripheral filament addition. This is further supported by earlier findings revealing that Zasp52, SALS, Fhos and DAAM, the four proteins also implicated in lateral sarcomere growth, likewise accumulate at the Z-disc (Bai et al., 2007; Farkas et al., 2024; Gonzalez-Morales et al., 2019; Molnar et al., 2014; Shwartz et al., 2016; Szikora et al., 2020a). Taking into account that (1) the myofibril thickening effect of FliIL674V depends on dLRRFIP, Fhos, and DAAM (and at least weakly on Zasp52), while the analogous effect of SALS depends on Fhos and DAAM (Farkas et al., 2024), and (2) all of these factors can be linked to actin regulation, we propose that they cooperate to promote the formation of a short, Z-disc-anchored actin filament that serves as a nucleation seed, or a precisely positioned “primer”, for subsequent actin elongation toward the H-zone (Fig. 7e). In this model the two formins, Fhos and DAAM would be required for the initial assembly of this short actin filament, while FliI/dLRRFIP would secure the stable integration of this filament. The roles of SALS and Zasp52 remain less clear, however, Zasp52 may contribute to the recruitment of α-Actinin, further enhancing filament integration through actin crosslinking, while SALS may act as a modulator within the barbed-end complex, where we envision dynamic regulation among CP, Fhos, DAAM and FliI/dLRRFIP. It is also worth noting that, although we found no *in vitro* evidence for formin regulation by *Drosophila* FliI (Pintér et al., 2020), human FliI has been shown to promote formin activation in cultured cells (Higashi et al., 2010), raising the possibility that FliI contributes to lateral myofibril growth by stimulating formin activity at the Z-disc, in addition to its critical role in filament integration. However, since the loss of FliI results in the accumulation of a large number of loosely associated myofilaments in the peripheral regions of myofibrils, clearly indicating that myofilament formation *per se* is not blocked, FliI is unlikely to play a critical role in Fhos and/or DAAM activation in this context.

An often overlooked but important aspect of sarcomeric structural organization is the precise alignment of filament ends at the Z-discs, A-I borders, and H-zone edges, a process achieved through largely unknown mechanisms. Based on our superresolution microscopy studies FliI is specifically enriched at the barbed end of the actin filaments, which is consistent with its *in vitro* ability to cap these ends (Pintér et al., 2020). Taking this together with its role in filament integration, it appears that the FliI/dLRRFIP complex may not simply promote lateral filament addition, but it does so in a precisely defined position (i.e. at the barbed ends), thereby providing a potential mechanism for placing barbed ends in register. Our data indicate that FliI/dLRRFIP is able to form higher order complexes, and may therefore act as an actin barbed-end crosslinker. Alternatively, a FliI/dLRRFIP trimer (or higher multimer) may associate with an as yet unidentified Z-disc protein through the dLRRFIP dimer, while simultaneously binding the barbed end of an actin filament through the GH domains of FliI.

Like FliI, the mammalian LRRFIP orthologs are best known as signaling proteins involved in pathways such as Wnt, TLR and NLRP3 (Dai et al., 2009; Gunawardena et al., 2011; Jin et al., 2013; Labbe et al., 2017; Liu and Yin, 1998). In addition, LRRFIP1 has been shown to be strongly expressed in the human and mouse heart and skeletal muscle (Labbe et al., 2017; Wei et al., 2015), and is required for myoblast differentiation in C2C12 cells (Wei et al., 2015). Mouse LRRFIP2 is also expressed in the heart and regulates embryonic cardiogenesis (Driss and Philippe Daubas, 2021). Although an uncharacterized RNAi phenotype had already linked the single *Drosophila* LRRFIP ortholog (CG8578) to muscle development (Schnorrer et al., 2010), the specific requirements during myofibrillogenesis uncovered here provide important novel insights into the mechanisms of this protein family. Remarkably, loss of FliI or dLRRFIP results in nearly identical IFM myofibrils phenotypes, and consistent with a concerted action, formation of the FliI/dLRRFIP complex is essential for myofibril formation. In contrast, the mouse knock-out phenotypes for FliI and LRRFIP2 are not identical to each other (Driss and Philippe Daubas, 2021; Kuwabara et al., 2023), but the early developmental defects and the seemingly complex roles in muscle growth and heart formation have precluded analysis of myofibrillogenesis at the relevant developmental stages in these mutants. Future investigations will therefore be needed to determine whether complex of the cardiomyopathy associated FliI protein and LRRFIP represents an evolutionary conserved module of peripheral myofilament integration during myofibril maturation.

## MATERIALS AND METHODS

### *Drosophila* stocks and genetics

All flies were raised and crossed at 25 °C according to standard procedures. *w^1118^* was used as the wild-type control. In addition, the following stocks were used: *y w; mef2-Gal4* (BDSC #27390), *fliI^3^* (BDSC #4730), *y^1^ fliI^sdby^* (Kyoto #101241), *y^1^ w* TI(CRIMIC.TG4.0)fliI^CR00462-TG4.0^*/FM7h (designated as *fliI^CRIMIC^*, BDSC #79264), *y^1^ v^1^*; *P(TRIP.JF02720)attP2* (designated as *fliI-KD*, BDSC #27566), *P(KK101150)VIE-260B* (designated as *dLRRFIP-KD^1^* VDRC v110146), *w^1118^, P(GD13973)v35968* (designated as *dLRRFIP-KD^2^*, VDRC v35968). For the viability assay third instar male larvae were collected with the proper genotypes and transferred to a fresh vial for aging. We have monitored the number of individuals survived till pupal and adult stage. For staging, white prepupae or newly eclosed flies were collected and kept until they reached the required age. In the actin incorporation assay, the *UAS-GFP::Act88F* transgene (BDSC #9253) (Roper et al., 2005) was expressed using the *fln-Gal4* driver (Bryantsev et al., 2012). The *fliI^3^* allele carries a point mutation in the GH2 domain causing an aminoacid replacement (G601S) (de Couet et al., 1995). The Kyoto #101241 stock, officially designated as *y^1^; fliI^8^* is also mentioned with the synonym as *fliI^sdby^*. However, based on former literature (de Couet et al., 1995; Deak et al., 1982) *fliI^8^* is a homozygous lethal allele that carries a 392 bps long deletion in the 5’ region of the gene (Campbell et al., 1993), whereas *fliI^sdby^* is a viable allele but no molecular information is available. Because the Kyoto #101241 stock is homozygous viable, we decided to clarify this discrepancy by DNA sequencing, and found that this *fliI* allele carries a single nucleotide change, causing an amino acid substitution in the GH6 domain (E1202K). Based on these findings this stock is neither homozygous lethal nor it carries the mutation described for *fliI^8^*, and for these reasons we conclude that it is certainly not a *fliI^8^*allele, instead it most likely corresponds to *fliI^sdby^*.

### Generation of FliI and dLRRFIP transgenes and mutant alleles

cDNAs encoding full-length FliI and dLRRFIP were cloned into the pENTR1a entry vector and used as templates for the generation of UAS-FliI and dLRRFIP transgenes. The N- and C-terminally tagged constructs were generated using Gateway cloning with the appropriate destination vectors (pPFW-attB and pPWF-attB for FliI; pTFW-attB and pTWF-attB for dLRRFIP). The FliI L674V mutant was generated by PCR-based site-directed mutagenesis using the pENTR1a-FL-FliI construct as a template. To generate the dLRRFIP binding surface mutants of FliI (B1, B2, and B1-2), the pENTR1a-FL-FliI vector was used as template. Two mutagenic oligonucleotides were designed, each spanning a portion of the coding sequence and introducing alanine substitutions at residues critical for binding in either surface 1 or surface 2. Using six primers, sequential PCR amplification followed by HiFi DNA assembly was performed to replace the corresponding regions in the template, yielding three constructs: B1 (surface 1 mutated), B2 (surface 2 mutated) and B1-2 (both surfaces mutated). All constructs were verified by DNA sequencing prior to transgenesis. Plasmids were injected into *Drosophila* embryos carrying the VIE-260B landing site (60100, VDRC) to generate transgenic lines. Primers used for construct generation are listed below.

The *dLRRFIP^K1^* mutant allele was generated by CRISPR/Cas9-mediated homology-directed repair, adapted from a previously described protocol (Yu et al., 2021). sgRNAs (5′-ATCCTTGTTATCACTTTCAA-3′ and 5′-TTGAAGGATCGCACCCGATG-3′) targeting the *dLRRFIP* locus were cloned into the pCFD5 vector. For the donor construct, 5′ and 3′ homology arms were PCR-amplified from genomic DNA of the *yw; CyO, vasa-long-gypsy-Cas9/Sco* strain and inserted into the pTV-GFP vector to generate a repair plasmid containing both arms. The repair plasmid and sgRNA constructs were co-injected into *yw; CyO, vasa-long-gypsy-Cas9/Sco* embryos. F_0_ adults were crossed to *w^1118^*, and targeted events were identified in the F_1_ generation based on GFP fluorescence in the larval CNS or in adult eyes. GFP positive individuals were selected, and deletion of the *dLRRFIP* locus was verified by sequencing.

For heterologous gene expression, cDNAs of FliI LRR and dLRRFIP and plasmid backbones (pET16b, pGEX, and pET16-eGFP) were amplified by Q5 polymerase (NEB, M0491S) followed by fragment isolation and HiFi assembly (NEB, E5520S) resulting the following constructs: pET16-(10xhis)dLRRFIP, pGEX-(GST)dLRRFIP, pET16-(10xhis)-eGFP-LRR. Plasmid sequences were confirmed by Sanger sequencing.

### Primer sequences

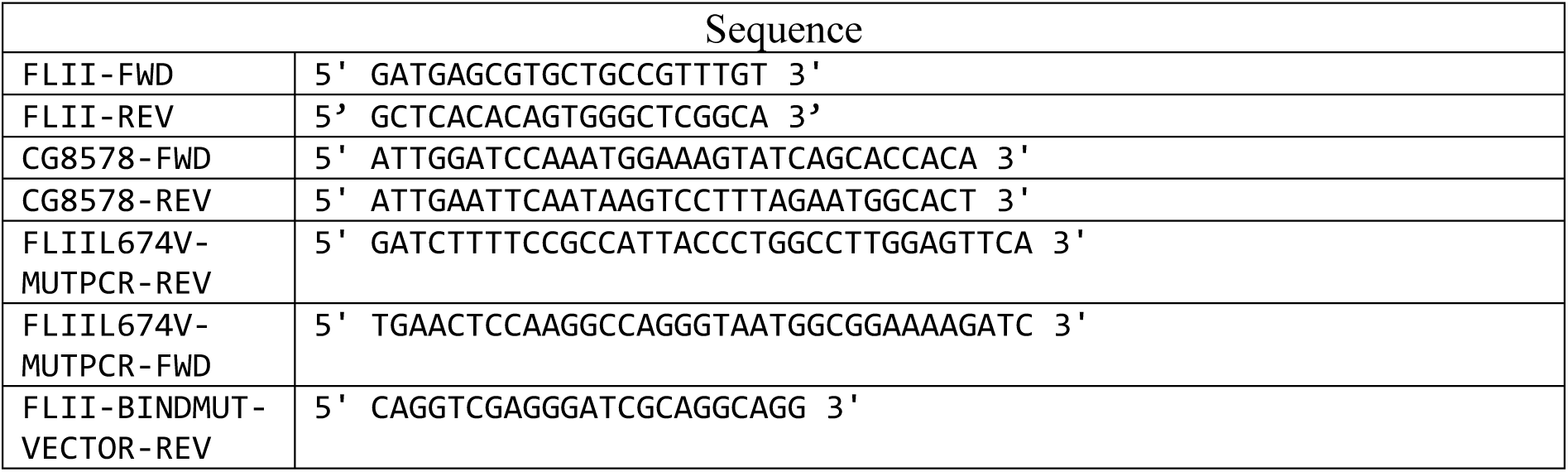
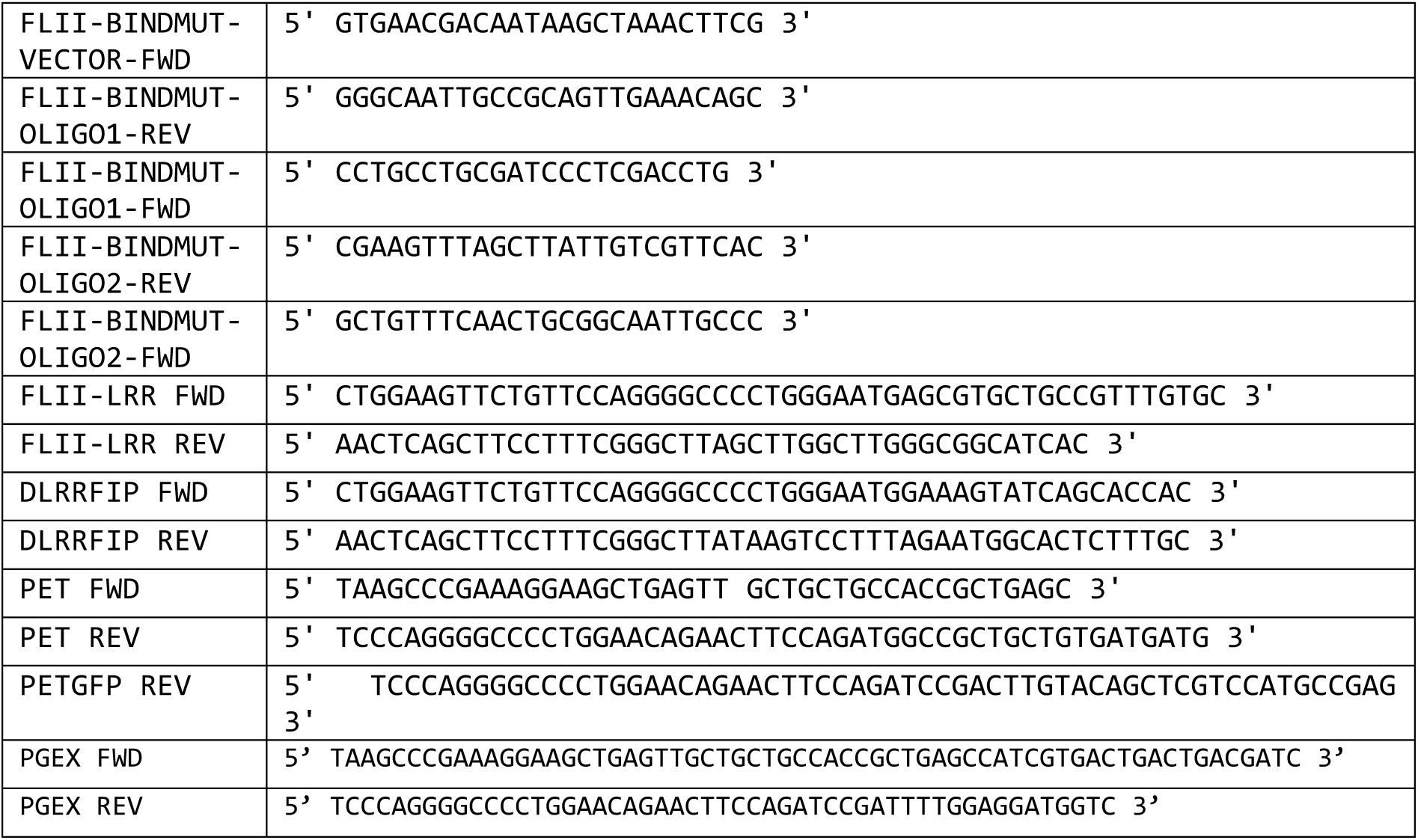

dLRRFIP B1 and B2 long oligos:

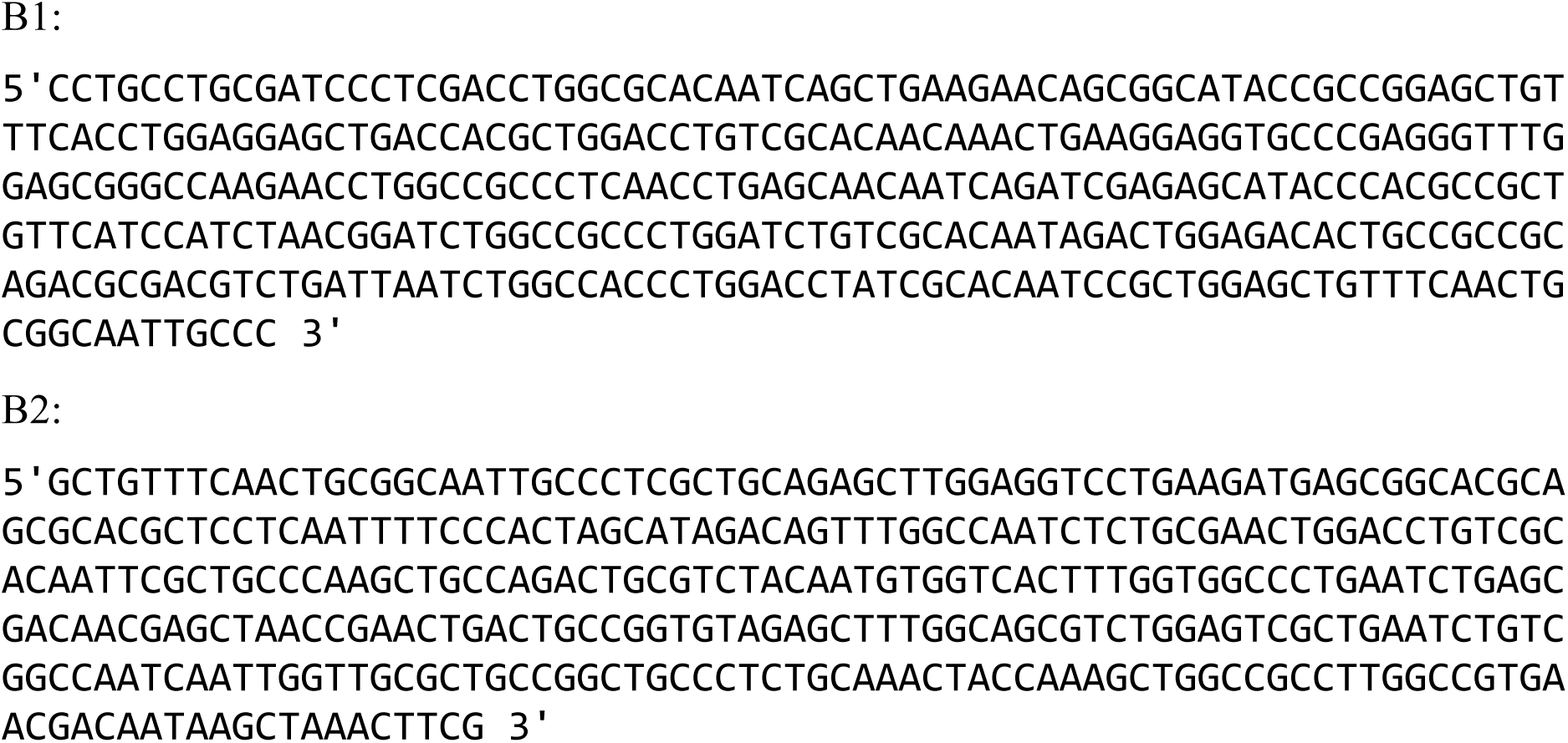

#### Flight assay

Flight tests were carried out with one day old flies. Flies in group of 5 were released in the middle of a perplex box illuminated from above. Based on their ability to fly, we divided them into 3 categories: fly up, horizontally or down. Those that flew upward or horizontally were counted as “flyer” and “weak flyer”, respectively, while the ones that fell downward as “non-flyer”.

### Sample preparation for fluorescent microscopy

#### IFM agarose sectioning

Thoraces were carefully dissected from the heads and abdomens using fine forceps, and the latter parts were discarded. The isolated thoraces were immediately transferred into a fixation solution containing 4% paraformaldehyde (Thermo Fisher Scientific; 16% stock) prepared in relaxing buffer (100 mM NaCl, 20 mM sodium phosphate, pH 7.0, 5 mM MgCl₂, 5 mM EGTA, and 5 mM ATP), and incubated overnight at 4 °C. Following a brief wash in PBS, the samples were embedded in warm 5% agarose (Lonza, SeaKem LE Agarose dissolved in PBS) and oriented as required. After the agarose solidified, the blocks were mounted onto a specimen holder and placed into a vibratome (Microm HM 650 V) filled with PBS. Sections of approximately 120 µm thickness were cut either parallel or perpendicular to the sagittal midline. The resulting sections were collected and stored in PBS at 4 °C until further processing. Standard immunohistochemistry protocols were subsequently performed on the sections.

#### IFM myofibril isolation

Individual myofibrils were obtained from the IFM following a modified version of the method described by Burkart et al. (Burkart et al., 2007). In brief, hemithoraces were first bisected and incubated on ice for 2 hours in relaxing solution supplemented with 50% glycerol. Subsequently, the DLM and/or DVM muscle fibers were carefully dissected and transferred into an Eppendorf tube, where they were dissociated by gentle pipetting in the presence of 0.5% Triton X-100. The resulting myofibril suspension was centrifuged at 10,000 × g for 2 minutes at 4 °C, after which the pellet was resuspended in 200 µl of 1× relaxing solution by pipetting. This washing step was repeated twice more. Following the final centrifugation, the fibers were again resuspended in relaxing solution. An aliquot of 20 µl was then applied onto a glass coverslip and fixed for 20 minutes using 4% paraformaldehyde (Thermo Fisher Scientific; 16% stock) in relaxing solution. See (Szikora et al., 2020b) for more details.

#### Thin filament isolation

Thin filaments were prepared from IFM using a protocol adapted from the thick filament isolation method described by Katzemich et al. (2012) (Katzemich et al., 2012). Briefly, individual myofibrils (isolated as outlined above) were subjected to limited Calpain digestion to detach thick filaments from the Z-disc (Contompasis et al., 2010; Reedy et al., 1981). The myofibrils were resuspended in 30 µl of buffer containing 20 mM imidazole (pH 6.8), 1 mM EDTA, 1 mM EGTA, 5 mM β-mercaptoethanol, and 30% glycerol, supplemented with 15 mM Calpain-1 (Merck). Calcium chloride was then added to a final concentration of 2 mM, and the mixture was incubated for 30 minutes at room temperature. Digestion was terminated by adding 100 µl of relaxing solution together with 100 mM Calpain Inhibitor I (Merck). The sample was subsequently passed five times through a 20G needle to mechanically separate the filaments, followed by centrifugation at 1500 × g for 3 minutes to pellet any remaining undigested material. The supernatant containing isolated filaments was applied to a glass coverslip and fixed for 20 minutes using 4% paraformaldehyde (Thermo Fisher Scientific; 16% stock). The fixed samples were then processed according to standard immunohistochemistry procedures.

#### Immunohistochemistry

Following fixation, samples were washed three times and incubated in PBS-BT blocking solution for 2 hours at room temperature (or 30 minutes in the case of isolated myofibrils). Primary antibodies diluted in blocking solution were applied and incubated overnight at 4 °C. The following primary antibodies were used: α-Actinin (DSHB 2G3-3D7, mouse, 1:100); Cpa (rabbit, 1:100; gift from Florence Janody) (Amandio et al., 2014); GFP (Abcam ab13970, chicken, 1:1000); Kettin Ig16 (DSHB BB17/29.5, rat, 1:200); Myosin 3E8 (DSHB 3E8-3D3, mouse, 1:1000); Myosin MAC147 (DSHB BB7/21.14, rat, 1:1000); Integrin-βPS (DSHB CF.6G11, mouse, 1:300); Obscurin Ig14–16 (rabbit, 1:400; gift from Belinda Bullard) (Burkart et al., 2007); FLAG (Merck M2, mouse, 1:1000). After washing, appropriate secondary antibodies were applied for 2 hours at room temperature. Detection was performed using highly cross-adsorbed goat anti-rabbit, anti-mouse, anti-rat, or anti-chicken IgG secondary antibodies conjugated to Alexa Fluor 405, 488, 546, or 647 (Life Technologies, 1:600). F-actin was visualized using Alexa Fluor 488-, or 546-conjugated phalloidin (Life Technologies), diluted in methanol (1:200). Nuclei were stained with DAPI (ThermoFisher Scientific, 1:500). After thorough washing, samples were mounted in ProLong Gold (P36930; Life Technologies) or stored in PBS prior to dSTORM imaging.

### Microscopy

#### Confocal microscopy

Images were acquired using a Zeiss LSM800 Airyscan confocal microscope with a Plan-Apochromat 63×/1.40 Oil DIC M27 or an EC-Plan-Neofluar 10×/0.30 M27 objective. Data were collected at Nyquist sampling.

#### dSTORM super-resolution imaging

Super-resolution imaging was performed as described previously (Szikora et al., 2020a; Szikora et al., 2020b). Experiments were carried out on a custom-built inverted microscope based on a Nikon Eclipse Ti-E platform, equipped with a Nikon CFI Apo 100× objective (NA = 1.49). Excitation was provided by a 647 nm laser (MPB Communications Inc., maximum power 300 mW), with the intensity at the sample adjusted to 2–4 kW/cm² using an acousto-optic tunable filter (AOTF). A 405 nm laser (Nichia, maximum power 60 mW) was employed to reactivate fluorophores. Fluorescence emission was recorded using an Andor iXon3 897 BV EMCCD camera (512 × 512 pixels, 16 µm pixel size), typically acquiring reduced field-of-view image stacks for dSTORM analysis. Excitation and emission light were separated using a multiband fluorescence filter set (Semrock, LF405/488/561/635-A-000) in combination with an additional emission filter (AHF, 690/70 H bandpass). Focus stability during acquisition was ensured by the microscope’s perfect focus system, keeping axial drift below 30 nm. Prior to imaging, the storage buffer was exchanged for GLOX switching buffer (van de Linde et al., 2011), and samples were mounted on microscope slides. Image sequences consisting of approximately 20,000–50,000 frames were collected with exposure times of 20–30 ms. Data analysis was performed using rainSTORM localization software (Rees et al., 2013). Single-molecule signals were fitted with a Gaussian point spread function to determine their centroid positions. Localization results were filtered based on intensity, localization precision (<20 nm), and point spread function width (0.8 ≤ σ ≤ 1.0). Sample drift due to thermal or mechanical factors was corrected using a correlation-based blind drift correction algorithm. See (Szikora et al., 2020b) for further details.

#### Transmission electron microscopy

Dissected hemithoraces, or whole pupae removed from their pupal cases and punctured in the abdomen with a fine needle, were fixed overnight at 4 °C in a solution containing 3.2% paraformaldehyde, 0.5% glutaraldehyde, 1% sucrose, and 0.028% CaCl₂ in 0.1 N sodium cacodylate buffer (pH 7.4). The samples were subsequently washed twice in the same buffer overnight at 4 °C, followed by a 15-minute rinse in distilled water. Post-fixation was carried out for 1 hour in 1% osmium tetroxide (Merck) prepared in distilled water. After osmium treatment, specimens were washed in distilled water for 10 minutes and dehydrated through a graded ethanol series (50% to 100%), with 10-minute incubations at each step and two changes in 100% ethanol. Dehydrated samples were then treated with propylene oxide (Molar Chemicals) for 5 minutes and embedded in an epoxy resin (Durcupan ACM; Sigma-Aldrich). Infiltration was performed using mixtures of propylene oxide and resin in ratios of 3:1, 1:1, and 1:3 (each for 1 hour), followed by two 1-hour incubations in pure resin and an additional overnight incubation at room temperature. Polymerization of the resin blocks was carried out at 56 °C for 48 hours. Following polymerization, blocks were trimmed and sectioned into 50 nm ultrathin slices using an Ultracut UCT ultramicrotome (Leica). Sections were collected on single-hole, formvar-coated copper grids (Electron Microscopy Sciences) and contrast-enhanced by staining with 2% uranyl acetate in 50% ethanol and 2% lead citrate in distilled water. Ultrathin sections were examined using a JEM-1400Flash transmission electron microscope (JEOL). Muscle sections were initially surveyed at low magnification (500–2000×), and representative images were acquired at the magnification of 12,000× for cross-sectional views and 8000× for longitudinal sections. Images were recorded as 16-bit grayscale files using a Matataki Flash 2k × 2k high-sensitivity sCMOS camera (JEOL) and saved in TIFF format.

### Image processing and analysis

#### Confocal image analysis

Sarcomere length and diameter were quantified directly from raw .czi files using a custom software tool, Individual Myofibril Analyser (Gorog et al., 2025) (https://github.com/GorogPeter94/Individual-Myofibril-Analyser-IMA-/tree/main). Isolated myofibrils were processed in automatic mode. Representative images were deconvolved using Huygens Professional Software (version 23.10; Scientific Volume Imaging). Optical sections were presented as average intensity projections, and brightness and contrast were adjusted linearly in Fiji (Schindelin et al., 2012).

#### TEM image analysis

For presentation purposes, image contrast was enhanced using Contrast Limited Adaptive Histogram Equalization (CLAHE) in Fiji (Schindelin et al., 2012). Center-to-center spacing between thick filaments was quantified following the approach described by Chakravorty et al. (2017) (Chakravorty et al., 2017). Measurements were performed on unprocessed TEM cross-sections in which filament orientation was properly aligned. A 512 × 512-pixel region was selected from a structurally intact area of each myofibril using the rectangular selection tool in ImageJ. The fast Fourier transform (FFT) of each region was then computed. In the resulting

FFT images, the distance from the center to the first-order diffraction peaks was determined by extracting intensity profiles along lines passing through the FFT origin. Gaussian functions were fitted to the peaks to locate their centers, and pixel distances between corresponding reflections were measured. These values were averaged to obtain d_pixels_avg. The real-space center-to-center spacing (S_real, nm) was calculated as follows: S_real = (pixel_size_original_nm × N_fft_pixels / d_pixels_avg) × (2/√3), where pixel_size_original_nm corresponds to the original TEM pixel size (nm), and N_fft_pixels denotes the FFT size (512).

#### dSTORM image analysis

Filtered and drift-corrected single-molecule localization datasets were analyzed as described previously (Erdelyi et al., 2015; Szikora et al., 2020a; Szikora et al., 2020b; Varga et al., 2023). Visualization and quantitative processing were performed using IFM Analyzer (version 2.1). For structural averaging, the built-in merge function was used to align localization datasets along the symmetry axes of the H-zone and I-band prior to averaging. Super-resolved images were then reconstructed from the merged localization lists using a pixel size of 10 nm. Quantitative measurements were performed on raw localization data derived from individual Z-disc regions.

### Biochemistry

#### Mass spectrometry

Whole thoraces (wild type and *fliI^CRIMIC.TG4^; UAS-FLAG::fliI-FL*) were flash-frozen in liquid nitrogen and homogenized in lysis buffer (50 mM Tris, 150 mM NaCl, 1% IGEPAL, 1 mM DTT, 3 mM pNpp, 1x protease inhibitor cocktail, 3 mM PMSF) using a TissueLyser (Qiagene). For immunopurification, 4 mg of total protein extract per sample was incubated with anti-DYKDDDDK antibody-coupled magnetic beads (30 μl, bead size: 50 nm, MACS® Technology, Miltenyi) following a modified protocol based on Hubner et al. 2010 and Lang et al. 2021 (Hubner et al., 2010; Lang et al., 2021). Following on-bead digestion with trypsin (Promega), samples were acidified and loaded onto single-use trapping mini-columns (Evotip; 1/8 of the total sample volume). Peptides were analyzed by data-dependent LC-MS/MS using an Evosep One system (88-min “15 SPD” built-in gradient) coupled online to an Orbitrap Fusion Lumos mass spectrometer (Thermo Fisher Scientific) operating in positive ion mode. MS1 spectra were acquired in the Orbitrap (R=60,000), while multiply charged precursor ions were selected based on cycle time (1.5 s) for HCD fragmentation and subsequent analysis in the ion trap.

Raw data were converted into peak lists using Proteome Discoverer (v 1.4) and searched against the Uniprot *Drosophila melanogaster* database, using the Protein Prospector search engine (v5.15.1) with the following parameters: enzyme: trypsin with maximum 2 missed cleavage; mass accuracies: 10 ppm for precursor ions and 0.6 Da for fragment ions (both monoisotopic); fixed modification: carbamidomethylation of Cys residues; variable modifications: acetylation of protein N-termini; Met oxidation; Met cleavage; cyclization of N-terminal Gln residues. Acceptance criteria: minimum scores: 22 and 15. 6 IPs (3 wild type, 3 FLAG-FliI samples) were used for meta analyses. The statistical analyzes were performed by edgeR (Robinson et al., 2010) using a modified spectral counting approach (PSMxCoverage %) (Branson and Freitas, 2016) to determine relative abundance of individual proteins (Lang et al., 2021). Thresholds for significance were set at a P-value < 0.01 and a minimum fold change of 1.5 relative to negative controls.

#### Co-immunoprecipitation, GST pull-down and Western blotting

A total of 2 × 10^6^ S2 cells per well were plated onto poly-L-Lysine-coated Labtech chamber slides and transfected with a total of 800 ng DNA using Effectene (Qiagen) according to the manufacturer’s instructions. 48 hours after transfection approximately 7×10^6^ S2 cells were harvested by centrifugation at 1000 x g for 2 minutes at 4°C. The cell pellet was then washed with 1 ml of cold PBS and re-centrifuged under the same conditions (1000 x g for 2 minutes at 4°C). Subsequently, the pellet was resuspended in 600 µl of FAC buffer (50 mM HEPES, 80 mM KCl, 5 mM MgCl2, 2 mM EGTA, 0.2% Triton X-100, pH 7.4), supplemented with a protease inhibitor mixture (Roche). The suspension was sonicated for 5 seconds to ensure complete lysis. Finally, cellular debris was removed by centrifugation at 12,000 x g for 10 minutes at 4°C. For each sample, 15 µl of anti-FLAG magnetic beads were used. The beads were then washed twice with 10 times their volume of TBS. After washing, the beads 600 µl of S2 cell lysate was added and incubated with gentle agitation for 2 hours at 4°C. Following incubation, the supernatant was carefully removed. The beads were then washed three times, each time with 1 ml of TBS, allowing the beads to remain in the wash buffer for 15 minutes per wash. Finally, the bound proteins were eluted by adding 40 µl of 1x SDS loading buffer (300 mM Tris-HCl, 10% SDS, 50% Glycerol, 25% β-mercaptoethanol, 0.05% bromophenol blue) to the beads and incubation at 100 °C for 4 minutes.

For GST pull-down experiments GST-tagged proteins were expressed in BL21 cells in LB media (10 g/l Tryptone, 10g/l NaCl, 5g/l Yeast extract) induced with 0.5 mM IPTG overnight at 18°C. Cells were sonicated in lysis buffer G (50mM Tris-HL pH 7.5, 4,65 mM DTT, 50 mM NaCl, 5 mM EDTA, 10% glycerol, 1x PIC) and cell debris was removed by centrifugation. Supernatant was subjected to Glutathione Sepharose 4B beads (Cytiva), equilibrated with lysis buffer and washed in wash buffer I (50 mM Trish-HCL pH 7.5, 5 mM DTT, 160 mM NaCl, 10% glycerol) and II (50mM Trish-HCL pH 7.5, 5 mM DTT, 50 mM NaCl, 5mM MgCl_2_, 10 mM KCl, 5% glycerol) and eluted in wash buffer II, supplemented with 50 mM reduced glutathione (pH7.6). The buffer was exchanged using D-10 Desalting Column (Merck) to storage buffer (50 mM HEPES pH7.6, 5mM DTT, 1.25% glycerol, 1% sucrose) and samples were concentrated with Amicon ultra 10k devices (Merck) and stored at −80°C. 15 ul of Glutathione Sepharose 4B beads were equilibrated in a storage buffer and incubated with purified GST-tagged protein, followed by washing. Then, S2 cell extracts containing the FLAG-tagged proteins were added to the beads and incubated for 3 hours at 4°C. Next beads were washed four times in buffer FAC (50 mM Hepes pH7.4, 80 mM KCl, 5 mM MgCl_2_, 2mM EGTA, 1% TritonX-100). For elution, 1xSDS loading buffer was added to the beads and incubated for 5 minutes at 100 °C.

Western blots were performed by using standard procedures. Mouse anti-FLAG (1:2000, M2, Sigma-Aldrich) and Mouse anti-Myc (1:1000, 9E10: sc-40 Santa Cruz Biotechnology) were used as primary antibodies. Anti-mouse IgG-HRP (light chain specific) (1:10000, 115-035-174 Jackson ImmunoResearch) was used as secondary antibody. Anti GST Tag clone DG122-2A7, HRP conjugate antibody (Merck 16-209) was also utilized in 1:2000 dilution. HRP conjugated proteins were visualized with the chemiluminescent Millipore Immobilon kit (Merck) using an Alliance Q9 Advanced chemiluminescence and epifluorescence imaging system (Uvitec).

#### Protein expression, purification, and size exclusion chromatography

Protein production was performed in Arctic express cells, DE3 (Agilent Technologies, Inc.) at 18 °C in 2YT auto-induction media (16 g/l tryptone, 10g/l yeast extract, 0.15 g/l MgSO_4_, 3.3 g/l (NH_4_)_2_SO_4_, 6.8 g/l KH_2_PO_4_, 7.1 g/l Na_2_HPO_4_, 0.5 g/l glucose, 2 g/l alpha lactose) for 48–60 hours. Cells were collected by centrifugation; if not utilized immediately, the cell pellets were preserved at −80 °C until needed. For lysis, cells were resuspended in buffer A (20 mM HEPES pH 7.5, 50 mM NaCl, 5% glycerol, 1% sucrose, 40 mM imidazole) supplemented with proteinase inhibitor cocktail (Roche). Both FliI LRR and dLRRFIP form inclusion bodies during heterologous expression, therefore buffer A was supplemented with 0.4% N-lauroylsarcosine (sarkosyl) as it was reported to facilitate protein solubilization (Tao et al., 2010). In the case of LRR we also used an N-terminal eGFP tag to facilitate its folding. The samples were sonicated for 20 cycles of 10 sec active sonication with 10 sec breaks at 40% amplitude, on ice to avoid overheating. The lysate was supplemented with 0.4% triton X-100 and centrifuged with 17000 g at 4 °C for 10 min. At this point the pellets still contains considerable amount of heterologous proteins, therefore the sonication process was repeated to increase the yield.

Both poly-histidine tagged eGFP-LRR and dLRRFIP are suitable for metal-affinity purification. Although GST-dLRRFIP does not bind to the metal affinity column by its own, its co-expression with 10xHis-dLRRFIP allows their co-purification. Cell extracts were loaded on nickel-NTA agarose column (ThermoScientific) and after washing with buffer A, heterologous proteins were eluted in buffer B (20 mM HEPES pH 7.5, 50 mM NaCl, 5% glycerol, 1% sucrose, 400 mM imidazole, 0.4% sarkosyl, 0.4% triton X-100). Buffer was exchanged to buffer GF (20 mM HEPES pH 7.5, 50 mM NaCl, 5% glycerol, 1% sucrose, 0.04% sarkosyl, 0.04% triton X-100) using PD MidiTrap G-25 (Cytiva) columns. Proteins were concentrated using Amicon ultra 10K filter units (Merck), then samples were flash frozen and stored at −80 °C or stored at 4 °C and used within a day.

Before gel filtration, samples were melted on ice and centrifuged at 17000 g for 10 minutes to remove aggregates and impurities. Gel filtration was performed on a Superdex 200 10/300 GL column (GE Healthcare) using 0.3ml/min flow rate, collecting 0.3 ml or 0.6 ml fractions starting at 7.5 ml from injection. The column was calibrated using the Broad range SDS-PAGE Standard (BioRad). The peaks of the 116.66 and 45 kDa proteins were marked on the related figures.

#### AlphaPulldown and structural modeling

To predict protein–protein interactions between FliI and dLRRFIP, we used the AlphaPulldown pipeline (Molodenskiy et al., 2025; Yu et al., 2023), which is based on AlphaFold2 (Jumper et al., 2021). This approach enables systematic prediction of protein–protein interactions by modelling defined domain combinations between a “bait” protein and a candidate interactor. Full-length and domain-segmented sequences of FliI and dLRRFIP were used to generate pairwise interaction models. FliI domains were treated as bait, while dLRRFIP served as the candidate protein. In total, 100 domain-to-domain combinations were evaluated, and five independent models were generated for each pair. Model quality was assessed using predicted Template Modelling (pTM) and interface pTM (ipTM) scores. Domain pairs with both pTM and ipTM scores exceeding 0.5 were considered for further analysis. Per-residue confidence was evaluated using predicted Local Distance Difference Test (pLDDT) scores. Based on these criteria, high-confidence models were selected for downstream structural interpretation. To further assess potential higher-order assemblies, trimeric models consisting of the FliI LRR domain and a dimeric form of dLRRFIP were generated using AlphaFold2 multimer, and models were filtered based on pTM, ipTM, and structural consistency. Protein–protein interfaces and interacting residues were identified using contact analysis in UCSF Chimera and interface evaluation with PDBePISA. Conserved interface residues were determined based on recurring contacts across high-confidence models. To assess the functional relevance of these residues, selected amino acids were subjected to *in silico* mutagenesis. The effects of these mutations on complex formation were evaluated by modelling trimeric FliI–dLRRFIP assemblies using AlphaFold3. Multiple mutant combinations were generated and compared to determine their impact on the stability and integrity of the predicted interaction interface. Among the tested variants, the minimal set of mutations required to disrupt the interaction (affecting 11 amino acids) was selected as the basis for the design of the B1, B2, and B1–2 constructs used in subsequent experimental analyses.

### Data analysis and figure preparation

Data were collected and organized using Microsoft Excel, while statistical analyses and data visualization were performed in GraphPad Prism 8 or R (R Core Team (2021)). For datasets comprising multiple independent experiments, superplots were used: individual data points represent pooled measurements to display distribution, variability, and sample size, whereas the mean of each independent experiment is plotted with error bars to reflect variability between experiments. Experiments were defined as independent when samples originated from separate parental crosses. Data normality was evaluated using either the D’Agostino–Pearson or Shapiro–Wilk tests. The specific statistical tests used for each analysis are indicated in the corresponding figure legends. A p-value of less than 0.05 was considered statistically significant. Part of the control data shown in figures 2**b**, 3**a**, 4**c** and 5**f** was already used in our Gorog et al., 2025 paper (Gorog et al., 2025). Final figures were prepared and assembled in Adobe Illustrator.

## ACKNOWLEDGEMENTS

We thank Belinda Bullard, Florance Janody, the Developmental Studies Hybridoma Bank, the Vienna Drosophila Resource Center, the Kyoto Drosophila Stock Center and the Bloomington *Drosophila* Stock Center for antibodies and fly stocks. We are indebted to Gyula Timinszky and Áron Szabó for critical reading and helpful comments on the manuscript. We thank Anikó Berente, Elvira Czvik, Anna Rehák, Bánfiné Erika Rácz and Velkeyné Ildikó Krausz for technical assistance.

