## Supplemental figures for "Flightless I and LRRFIP work together to regulate lateral growth of the sarcomeres in *Drosophila*"

### SUPPLEMENTARY INFORMATION

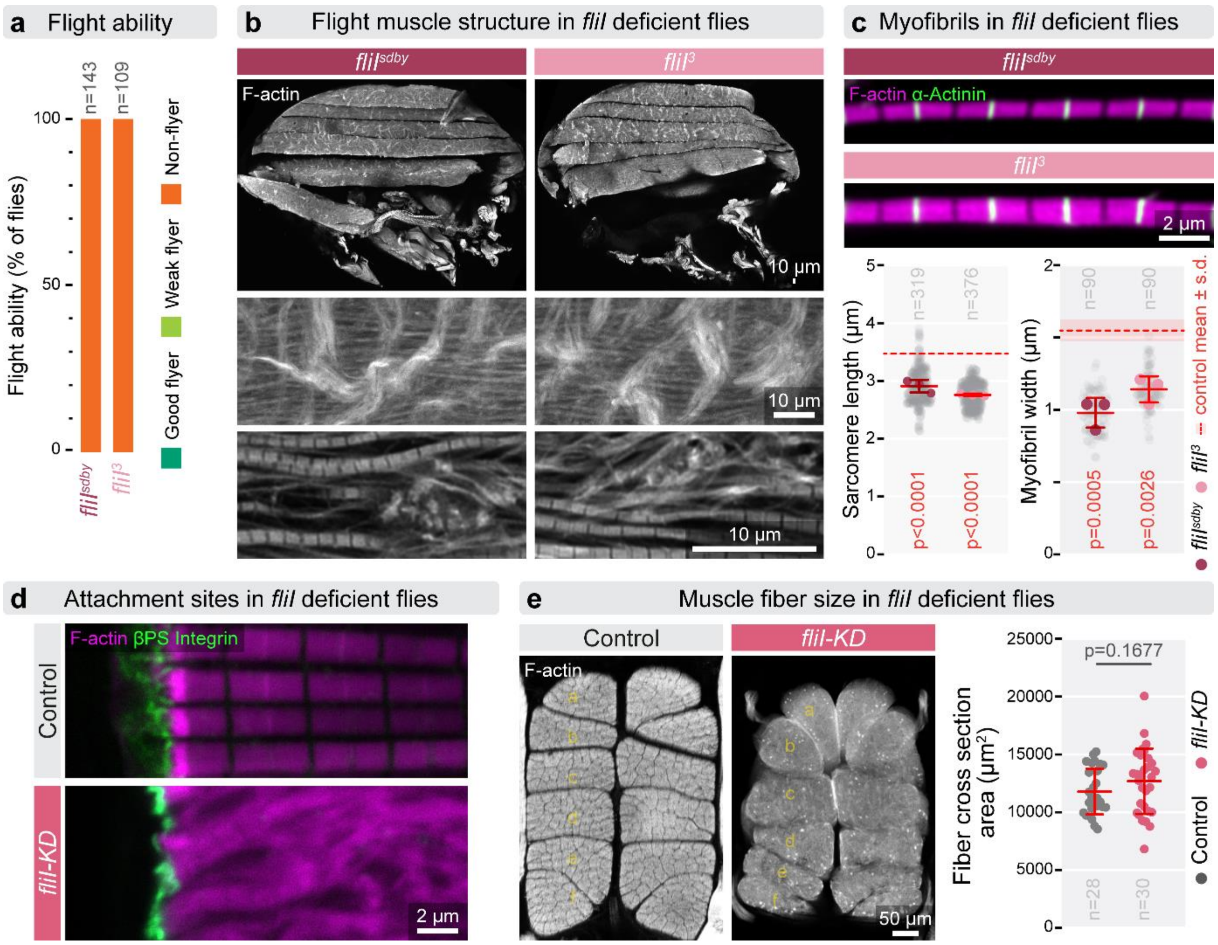


**Supplementary Figure 1. The IFM defects caused by hypomorphic *fliI* alleles, and muscle fiber organization upon FliI knockdown. (a)** Flight assay showing complete loss of flight in $fliI^{sdby}$ *and* $fli^{3}$ mutant flies at ~24 hours AE. **(b)** Low- and high-magnification images of thoracic sections revealing DLM fiber structure. Both in $fliI^{sdby}$ and $fli^{3}$ animals, large myofibrillar aggregates are visible on the surface of otherwise normally sized fibers. At higher resolution, individual myofibrils appear thinner, disorganized, and exhibit severely disrupted sarcomeric structure with loss of the regular actin pattern. F-actin is visualized by phalloidin staining (gray). **(c)** Top panels show isolated myofibrils from $fliI^{sdby}$ *and* $fli^{3}$ IFMs, demonstrating reduced sarcomere length and myofibril diameter in the hypomorphic mutant animals. The graphs below show morphometric quantifications, confirming significant reductions in both parameters. Statistical significance was assessed using one-way ANOVA followed by Dunnett's multiple-comparisons test. Light gray dots represent individual measurements, while larger dots indicate mean values from independent experiments. Error bars show mean of independent experiments ± s.d., and n denotes the number of independent measurements. The mean ± s.d. of control ($w^{1118}$) is indicated. (**d**) Confocal images of IFM attachment sites in control and *fliI-KD* (*mef2-Gal4/UAS-TRIP.JF02720*) muscles. Although myofibrils are disorganized, attachment sites remain intact in *fliI-KD* muscles. F-actin is labeled with phalloidin (magenta), and attachment sites are marked by βPS-integrin (green). (**e**) Left: thoracic cross-sections showing DLM fibers in control and *fliI-KD* animals, indicating normal fiber number. Right: quantification of DLM cross-sectional area shows comparable fiber size between control and *fliI-KD* IFMs. Statistical significance was assessed using unpaired t-tests. Raw data are provided in SFig1SourceData.

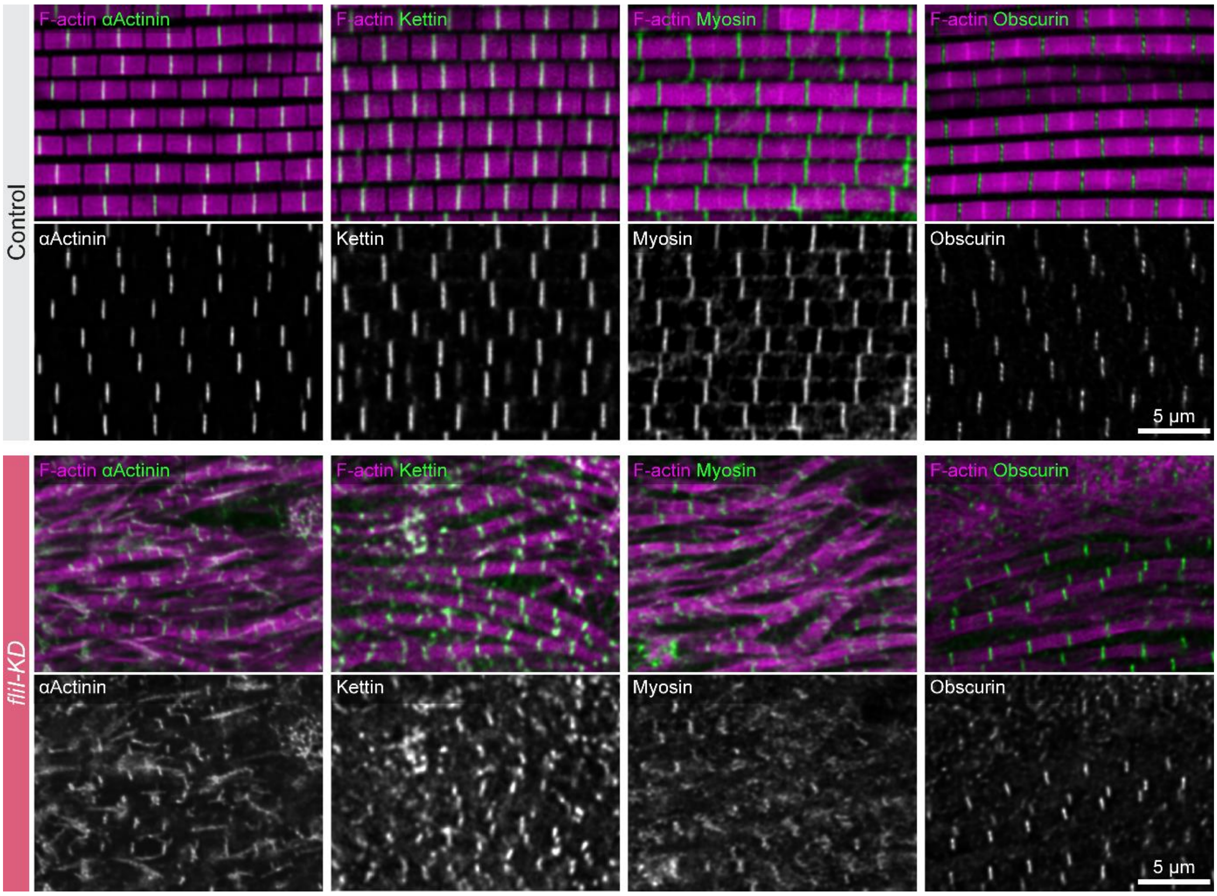


**Supplementary Figure 2. Distribution of the major sarcomeric markers upon FliI knockdown.** Confocal images of IFM myofibers from control (upper rows) and *FliI-KD* (*mef2-Gal4/UAS-TRIP.JF02720*) (lower rows) hemi-thoraces stained for actin (in magenta) as a thin filament marker, α-Actinin as a Z-disc marker, Kettin as an elastic filament (and Z-disc) marker, Myosin as a thick filament marker, and Obscurin as an M-line/H-zone marker (all in green in the overlay images, and in grey in the single channel images). Contrasting to the highly regular distribution pattern of these proteins in the controls, the *FliI-KD* muscles exhibit a partial loss of normal myofibril organization, yet they contain thin myofibrils with irregularities in their diameter, actin, α-Actinin, Kettin and Myosin accumulation patterns, while displaying largely normal looking Obscurin enrichment in the H-zone. Raw data are provided in SFig2SourceData.

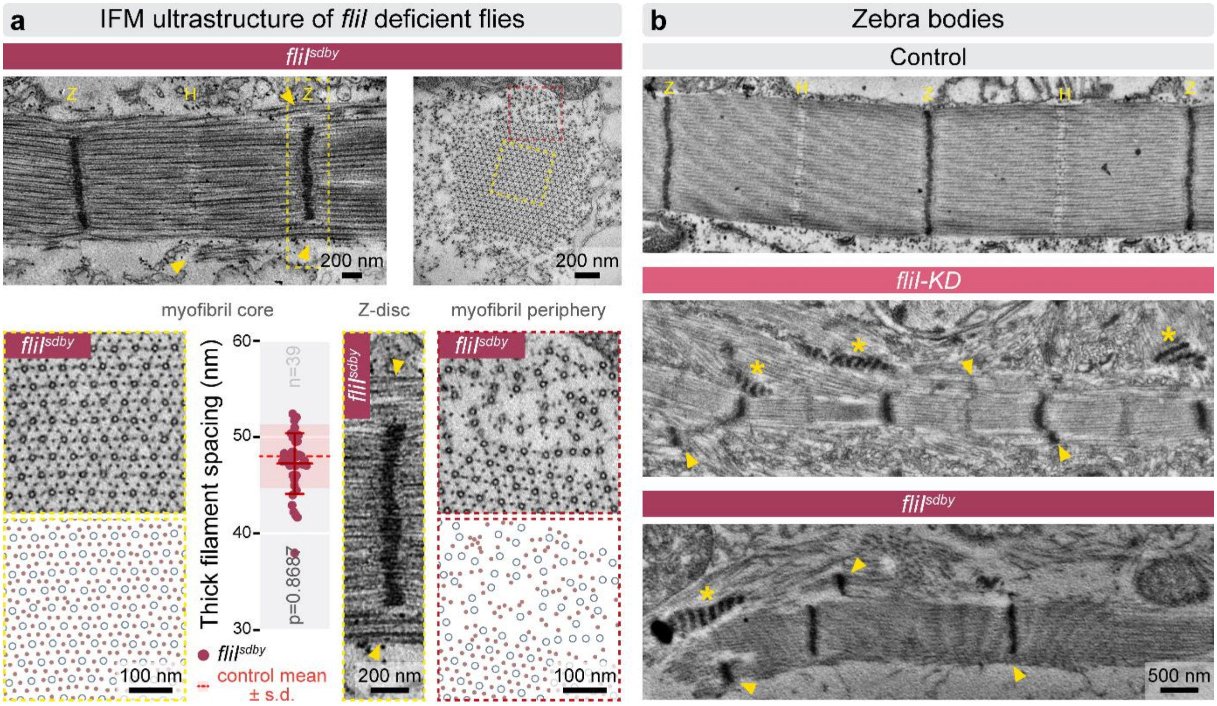


**Supplementary Figure 3. TEM analysis of *fliI* deficient myofibrils with Z-disc defects and zebra bodies.** (**a**) Electron micrographs of longitudinal and cross-sections of IFM myofibrils show that sarcomeres in *fliI*$^{sdby}$ mutants are shorter and thinner than in controls. Z-discs (yellow "Z") frequently fail to span the entire width of the myofibrils and are instead restricted to the core region, leaving peripheral myofilaments partially or completely unintegrated (yellow arrowheads). Boxed regions (dotted rectangles), shown at higher magnification below, reveal that in *fliI*$^{sdby}$ mutants the myofibrils are surrounded by unincorporated thin and thick filaments (boxes on the right), while their myofibril core (boxes on the left) exhibits the characteristic hexagonal packing. The Z-discs are smaller than normal, and often fails to extend to the myofibril periphery (yellow arrowheads), although Z-disc width and organization within the core remain comparable to controls. Despite these defects, the core myofilament lattice retains normal hexagonal organization and spacing, as indicated by unchanged average thick filament distances ($p = 0.8687$). Statistical significance was assessed using a Mann-Whitney test. Error bars represent mean ± s.d.; n indicates independent measurements. Mean ± s.d. of control (*w*$^{1118}$) is indicated. (**b**) In addition to Z-disc defects and impaired peripheral myofilament integration (yellow arrowheads), prominent zebra body–like, electron-dense myofibrillar aggregates (yellow asterisks) are observed in both *fliI-KD* and *fliI*$^{sdby}$ IFMs. Raw data are provided in SFig3SourceData.

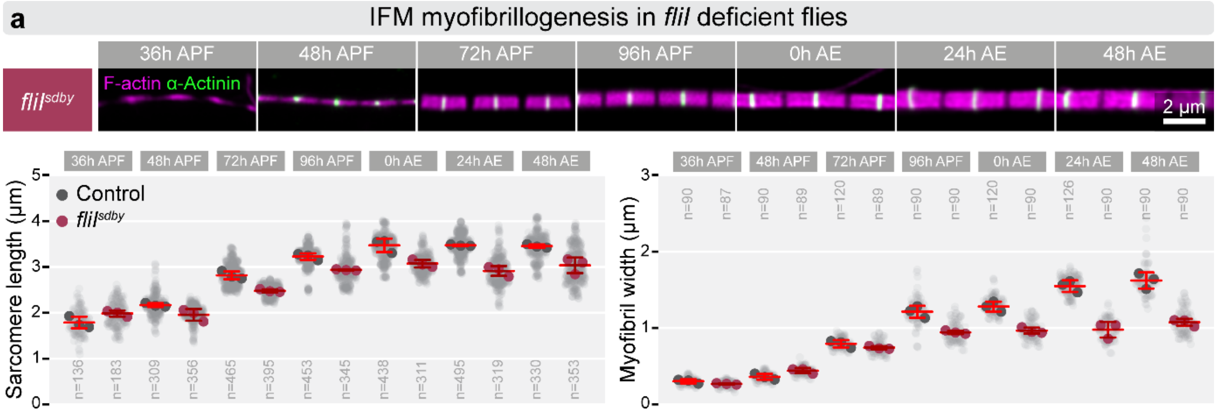


**Supplementary Figure 4. Developmental analysis of sarcomere length and width in *fliI*$^{sdby}$ mutant myofibrils.** (**a**) Representative images of isolated myofibrils show sarcomere growth in *fliI*$^{sdby}$ IFMs across seven time points (from 36 hours APF to 48 hours AE). F-actin (magenta) is labeled with phalloidin, and Z-discs (green) with α-Actinin. Graphs below show quantification of sarcomere length and myofibril width during development. Light gray dots represent mean values for individual measurements, and larger dots indicate means from independent experiments. The mean and s.d. of these experiments are provided, and n denotes the number of independent measurements. At 36 and 48 hours APF, no differences are observed between control and *fliI*$^{sdby}$. From 72 hours APF onward, sarcomere length is significantly reduced in *fliI*$^{sdby}$ IFMs ($p<0.0001$). Myofibril width is initially comparable but becomes significantly reduced from 96 hours APF through adulthood ($p<0.0001$). Statistical significance was assessed using two-way ANOVA followed by Dunnett's multiple-comparison test. Raw data are provided in SFig4SourceData.

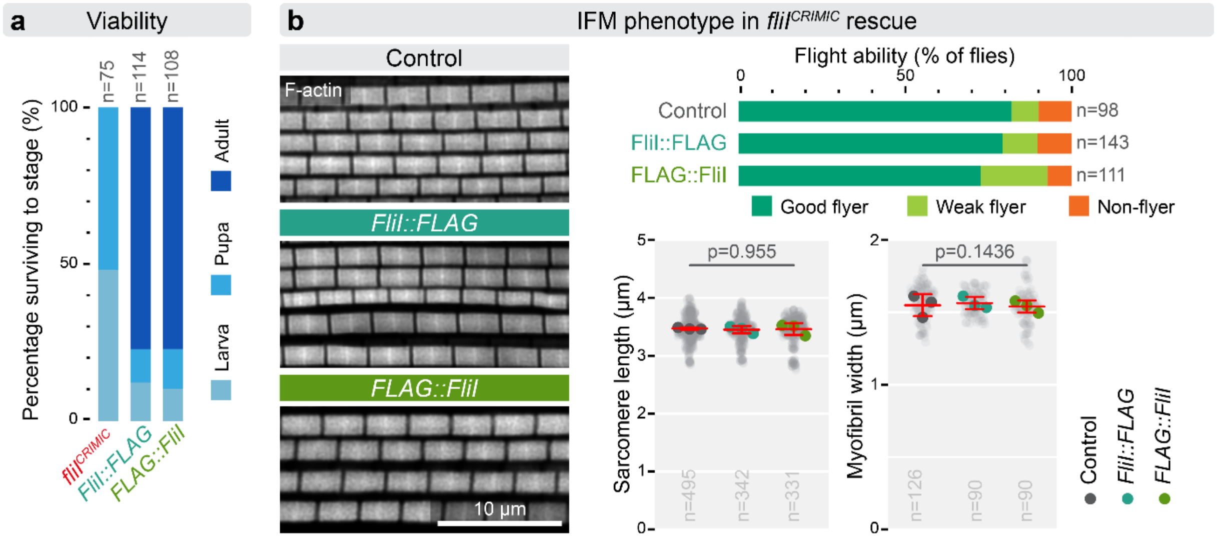


**Supplementary Figure 5. Rescue of the *fliI*$^{CRIMIC}$ null phenotype by FLAG tagged full length FliI.** (**a**) Graph showing viability (% survival) of flies in a *fliI*$^{CRIMIC}$ null mutant background, with or without rescue by N- or C-terminally FLAG-tagged FliI, expressed under the control of CRIMIC-Gal4. *n* indicates the number of larvae tested. (**b**) Left: high-resolution images of IFM myofibrils from control and *fliI* rescue animals, showing normal muscle structure upon rescue. F-actin is visualized by phalloidin staining (gray). Right (top): flight assay demonstrating full restoration of flight ability in both N- and C-terminal FLAG-tagged rescue lines. The graphs below show morphometric quantification of sarcomere size, confirming complete rescue of both length and width (p=0.955 and 0.1436, respectively). Statistical significance was assessed using one-way ANOVA followed by Dunnett's multiple-comparison test. Light gray dots represent individual measurements, while larger dots indicate mean values from independent experiments. Error bars show mean of independent experiments ± s.d., and n denotes the number of independent measurements. Raw data are provided in SFig5SourceData.

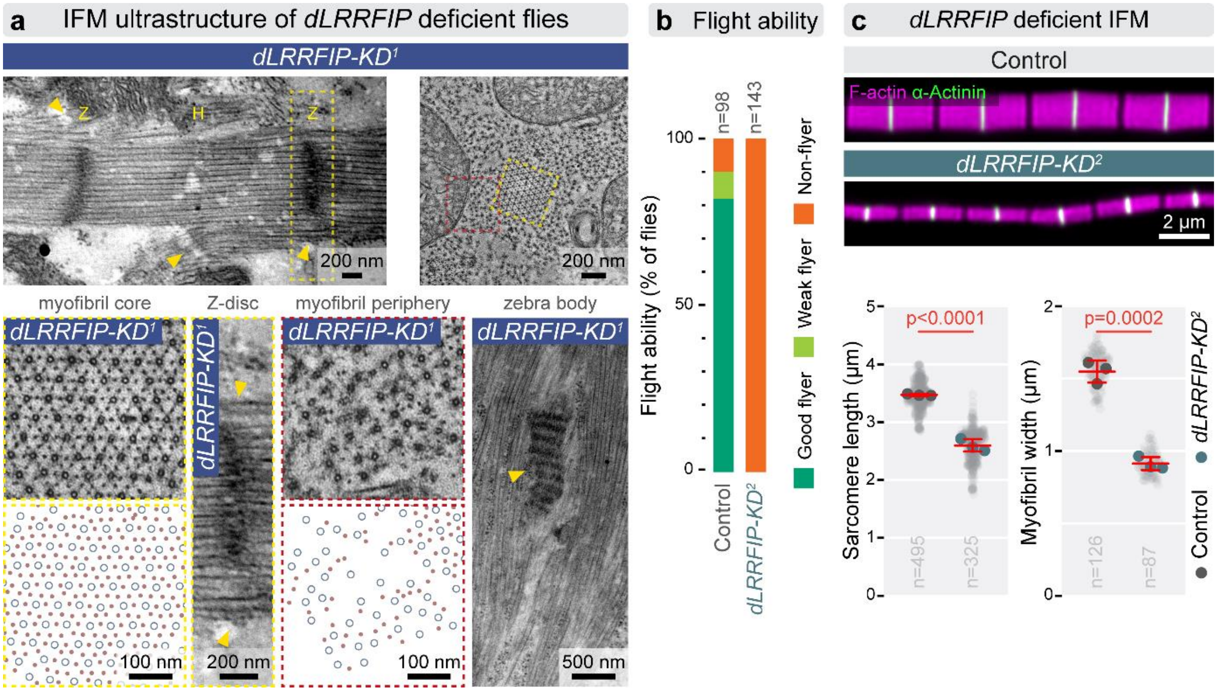


**Supplementary Figure 6. The loss of dLRRFIP affects peripheral myofilament integration and Z-discs formation, resulting in myofibril growth defects.** (**a**) Electron micrographs of longitudinal and cross-sections of IFM myofibrils show that sarcomeres in *dLRRFIP-KD*$^{1}$ (*mef2-Gal4/UAS-KK101150*) IFM are shorter and thinner than in controls. Z-discs (yellow "Z") are often smaller than normal and do not span the entire width of the myofibrils, leaving peripheral myofilaments partially or completely unintegrated (yellow arrowheads). Boxed regions (dotted rectangles), shown at higher magnification below, reveal myofibrils surrounded by unincorporated thin and thick filaments that lack the characteristic hexagonal packing in the peripheral region (dark pink boxes), while exhibiting a wild type pattern in the core region (yellow boxes). In between the boxes, an enlarged Z-disc, confined to the myofibril core, is also shown. In addition, prominent, zebra body–like, electron-dense myofibrillar aggregates are observed (yellow arrowhead) in *dLRRFIP-KD*$^{1}$ IFMs. (**b**) Quantification of the flight ability of control ($w^{1118}$) and *dLRRFIP-KD*$^{2}$ flies 24 hours AE. (**c**) Top panels show isolated myofibrils from control and *dLRRFIP-KD*$^{2}$ (*mef2-Gal4/UAS-GD13973*) IFMs, demonstrating reduced sarcomere length and myofibril diameter in knockdown animals. The graphs below show quantification of these parameters, confirming a significant reduction in both of them ($p<0.0001$ and $p=0.0002$, respectively). Statistical significance was assessed using unpaired t-tests. Light gray dots represent individual measurements, while larger dots indicate mean values from independent experiments. Error bars show mean of independent experiments ± s.d., and n denotes the number of independent measurements. Raw data are provided in SFig6SourceData.

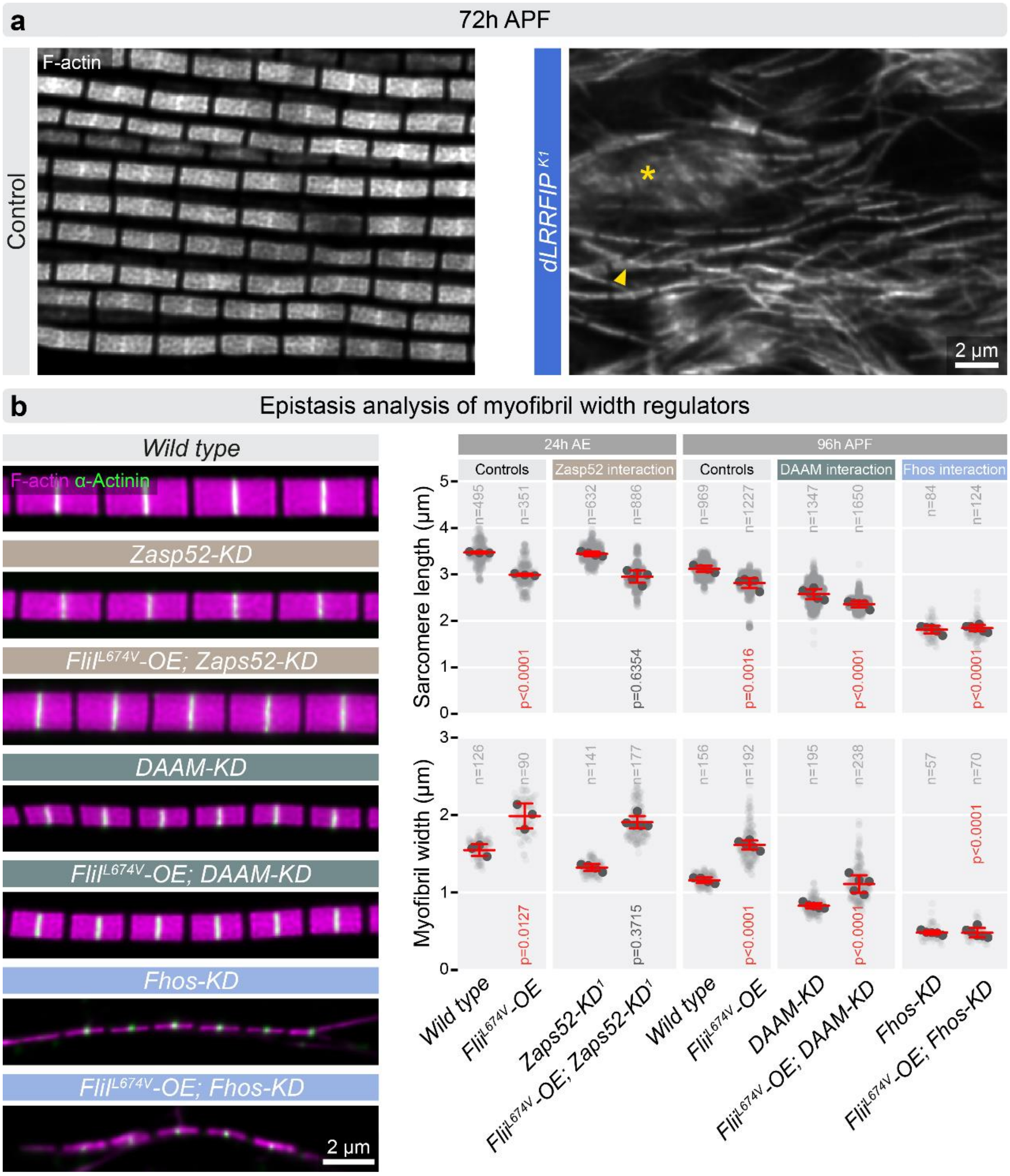


**Supplementary Figure 7. IFM phenotype of the *dLRRFIP* null mutant, and genetic interaction assays between the myofibril width regulators.** (**a**) F-actin staining of a pupal IFM muscle fiber from control (left) and *dLRRFIP*$^{K1}$ null mutant (right), 72 hours APF. Note the thin myofibrils (arrowhead) in the mutant, together with the presence of actin aggregates (asterisk). (**b**) Individual myofibrils stained for actin (magenta) and α-Actinin (green) to mark the Z-discs from IFMs of the genotype indicated. Note that overexpression (OE) of FliI$^{L674V}$ strongly increases myofibril width, while the knockdown of Fhos, DAAM and Zasp52 reduces myofibril diameter (with a various efficiency, Fhos exhibiting the strongest effect). Light gray dots represent individual measurements, while larger dots indicate mean values from independent experiments. Error bars show mean of independent experiments ± s.d., and n denotes the number of independent measurements. Statistical significance was assessed either using unpaired t-tests or One-way ANOVA with Dunnett's multiple comparison. Raw data are provided in SFig7SourceData.

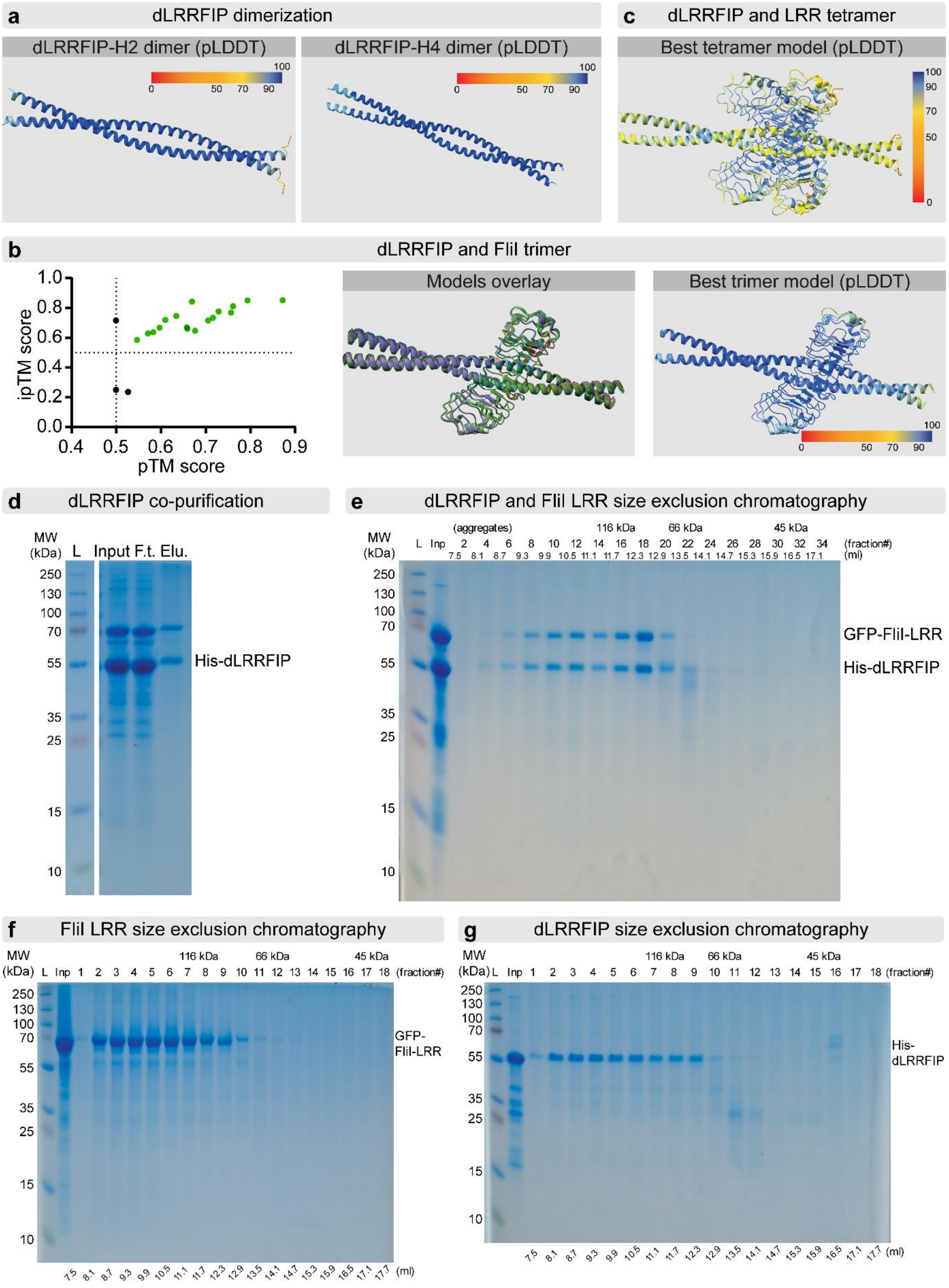


**Supplementary Figure 8. Interactions between FliI LRR and dLRRFIP revealed by *in silico* predictions and biochemical analysis.** (**a**) AlphaFold2 models of the dimerization of the H2 and H4 helices of dLRRFIP. Colors indicate the pLDDT score: dark blue signifies predictions with high confidence, red signifies low confidence. (**b**) AlphaPulldown analysis indicates the formation of a trimeric FliI-LRR/dLRRFIP complex. All the models above the cut-off of 0.5 pTM and iPTM scores are similar, and the pLDDT scores also indicate a confident prediction. (**c**) The predicted tetrameric structure of dLRRFIP and FliI-LRR, colored by pLDDT scores. (**d, e**) Uncropped images of polyacrylamide gels shown in Figure 7B and 7C, respectively. GFP-LRR (**f**) and His-dLRRFIP (**g**) were purified by metal affinity and gel-filtrated on Superdex 200 10/300 GL column. Both proteins appear in fractions near to the exclusion volume (7.5ml) indicating the formation of polymers or aggregates. Note that FliI LRR is present in fractions where the monomeric form (73 kDa) would be expected, while dLRRFIP can only be detected in fractions where the dimeric or the higher order structures would be expected.

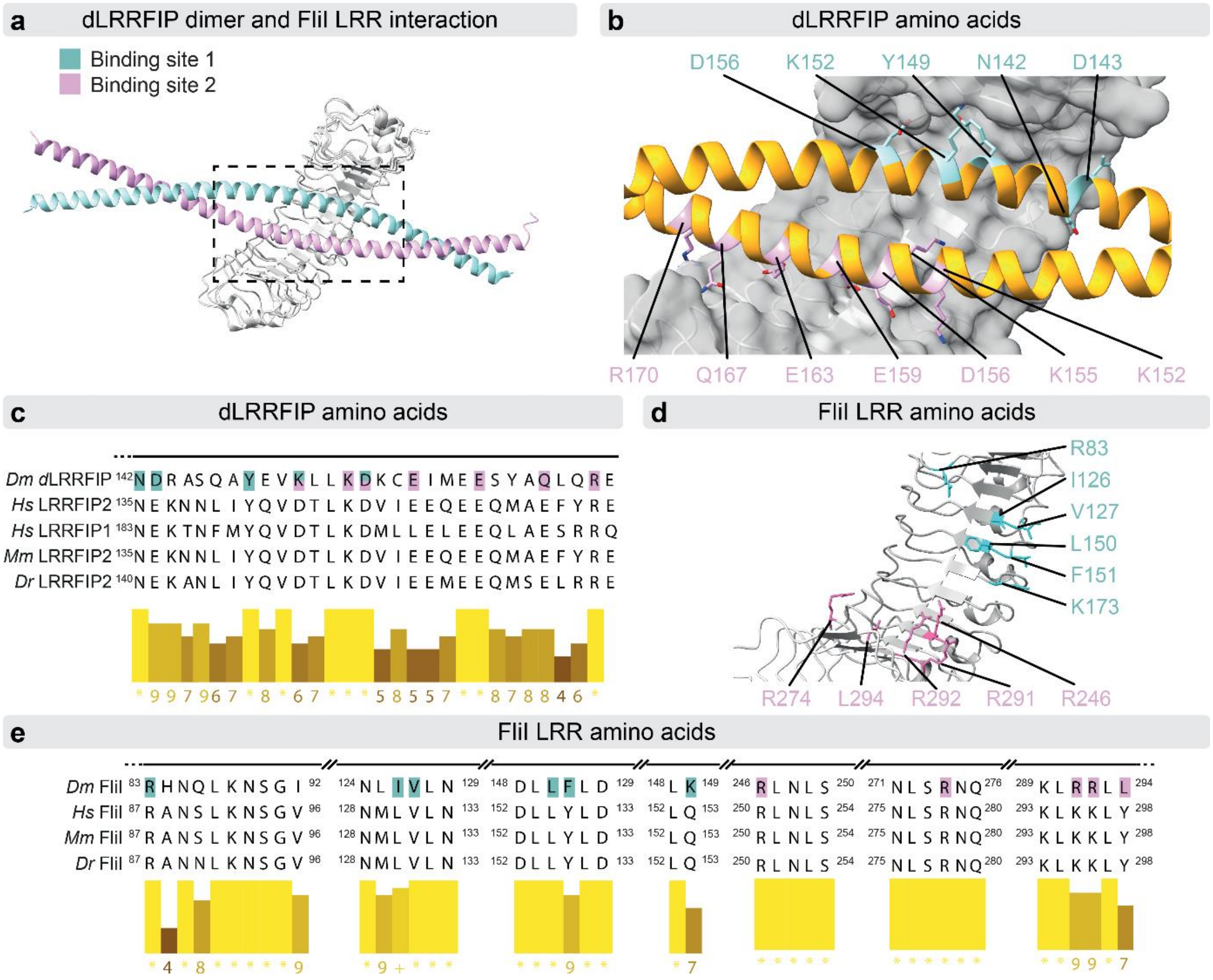


**Supplementary Figure 9. Structural prediction for the dLRRFIP-FliI LRR interaction.** (**a**) dLRRFIP interacts with the FliI LRR domain in a dimeric form, and both proteins possess two major interaction surfaces. (**b**) Amino acids of dLRRFIP that potentially interact with FliI-LRR are shown in a molecular model, and they are also highlighted in a multiple sequence alignment in (**c**). (**d**) Amino acids of FliI-LRR that potentially interact with dLRRFIP are shown in a molecule model, as well as in a multiple sequence alignment below in (**e**). To create the FliI LRR B1, B2 and B1-2 mutants these amino acids were mutated to Alanin. Cyan indicates the residues of B1, while pink shows the B2 related amino acids. *Dm: Drosophila melanogaster, Hs: Homo sapiens, Mm: Mus musculus Dr: Danio rerio.*
